# To make a short story long: simultaneous short and long RNA profiling on Nanopore devices

**DOI:** 10.1101/2022.12.16.520507

**Authors:** Morgan MacKenzie, Susan Tigert, Debbie Lovato, Hamza Mir, Kamyar Zahedi, Sharon L. Barone, Marybeth Brooks, Manoocher Soleimani, Christos Argyropoulos

## Abstract

Sequencing of long coding RNAs informs about the abundance and the novelty in the transcriptome, while sequencing of short coding RNAs (e.g., microRNAs) or long non-coding RNAs informs about the epigenetic regulation of the transcriptome. Currently, each of these goals is addressed by separate sequencing experiments given the different physical characteristics of RNA species from biological samples. Sequencing of both short and long RNAs from the same experimental run has not been reported for long-read Nanopore sequencing to date and only recently has been achieved for short-read (Illumina) methods. We propose a library preparation method capable of simultaneously profiling short and long RNA reads in the same library on the Nanopore platform and provide the relevant bioinformatics workflows to support the goals of RNA quantification. Using a variety of synthetic samples we demonstrate that the proposed method can simultaneously detect short and long RNAs in a manner that is linear over 5 orders of magnitude for RNA abundance and three orders of magnitude for RNA length. In biological samples the proposed method is capable of profiling a wider variety of short and long non-coding RNAs when compared against the existing Smart-seq protocols for Illumina and Nanopore sequencing.

## Introduction

Long coding and noncoding (short or long >200 nt long) RNAs yield valuable information about the abundance and novelty of the transcriptome and its epigenetic regulation respectively. Noncoding RNAs are of interest for clinical research applications, as their relative stability and tissue-specific nature make them viable candidates for disease-state biomarkers (1–6). There is currently a need to simultaneously sequence coding and non-coding RNAs from the same sample in a convenient and robust manner. In particular, consideration of epigenetic regulation requires examination of the quantitative relationships between noncoding and coding RNAs or between categories of noncoding RNAs e.g. microRNAs and long noncoding (lncRNA) RNAs (7–11). On the biomarker side, there exist multiple proposals for cell-free RNA based panels derived from either microRNAs(2, 4, 12, 13) or lncRNAs(14, 15), but there has been little work to combine markers from both categories, or even coding RNAs(16) and evaluate them in a prospective rigorous manner. This is in no small part due to the biochemical incompatibility of the existing sequencing protocols for non-coding and coding RNAs, which require construction of separate libraries that are then sequenced in parallel(17).

Approaches to simultaneously sequence RNAs from multiple classes like Holo-Seq(18) and Smart-seq-total(19) target short-read sequencing platforms. In recent years, long-read platforms(20) such as those by Oxford Nanopore Technologies (ONT) that span the entire range from portable devices to large scale high throughput sequencers have emerged as an alternative to short read sequencing. While the spectrum of applications of Nanopore sequencing is extremely wide, ranging from genomic sequencing to epigenomics and transcriptomics, there currently does not exist a method to simultaneously profile short and long RNAs in this platform. In fact, most library protocols for Nanopore sequencing exclude cDNAs derived from short RNAs. This represents a significant missed opportunity, because Nanopore sequencing provides the most accessible platform, in terms of acquisition, maintenance and operational costs, with a portability profile that is unmatched by all other alternatives.

We report here the first simultaneous sequencing of short and long RNAs on Nanopore devices using a single tube modification of ONT’s rapid PCR-cDNA (SQK-PCS109) library preparation kit. This approach polyadenylates all RNAs in a sample as a means for making all species compatible for use in SMART-Seq protocols. We demonstrate that this modification can be used to quantitate miRNAs and long-coding RNAs in synthetic mixes and generate counts that are in direct proportion to the abundance of RNAs in the sample. Using biological samples derived from homogenized tissues, we show that the proposed method delivers sequencing profiles that differ from those generated by Illumina sequencing and ONT’s unmodified protocols. The data and analyses presented herein go a long way to deliver a sequencing workflow and its bioinformatics ecosystem that allows for the simultaneous analysis of coding and noncoding RNAs for research applications.

## Materials and Methods

### Poly-Adenylation Lengthening of Short RNAs for Simultaneous Short and Long Nanopore Sequencing (PALS-NS)

#### Biochemical workflow/library preparation

Our proposed protocol for the simultaneous detection of short and long RNAs polyadenylates all RNAs in a sample before using them as input to any Smart-seq protocol(21) for long sequences (such as Oxford Nanopore’s SQK-PCS109) that requires polyadenylated, poly(A)+, RNA. The major change introduced is the execution of the poly-adenylation in the same tube as the reverse transcription (RT) and template switching reactions (**Figure *1***A), similar to the (Capture and Amplification by Tailing and Switching, CATS(22)/D-Plex Small RNA-seq(23)) and Smart-seq-total (19) workflows for Illumina sequencing. The remaining steps of the Smart-seq protocol are carried out without modifications: reverse transcription (RT) using a poly-T, VNP primer, addition of non-templated nucleotides (usually cytidines) strand switching via a strand switching primer (SSP) that contains a short ribo-nucleotide tail and PCR using universal primers that amplify between the 5’ end of the SSP and the 3’end of the VNP. Finally, the amplified library is purified using AMPure XP (or equivalent) beads, the rapid sequencing adaptors are added, and the sample is loaded on the flow cell for sequencing (**Figure *1***A). The brief duration of the poly-A tailing reaction in PALS-NS requires one further change to the Nanopore RNAseq protocols: size selection should be performed with 1.8x volumetric ratio of beads to library in order to retain both short and long cDNAs, instead of the usual 0.8x-1.0x ratio for long read sequencing.

**Figure 1.**
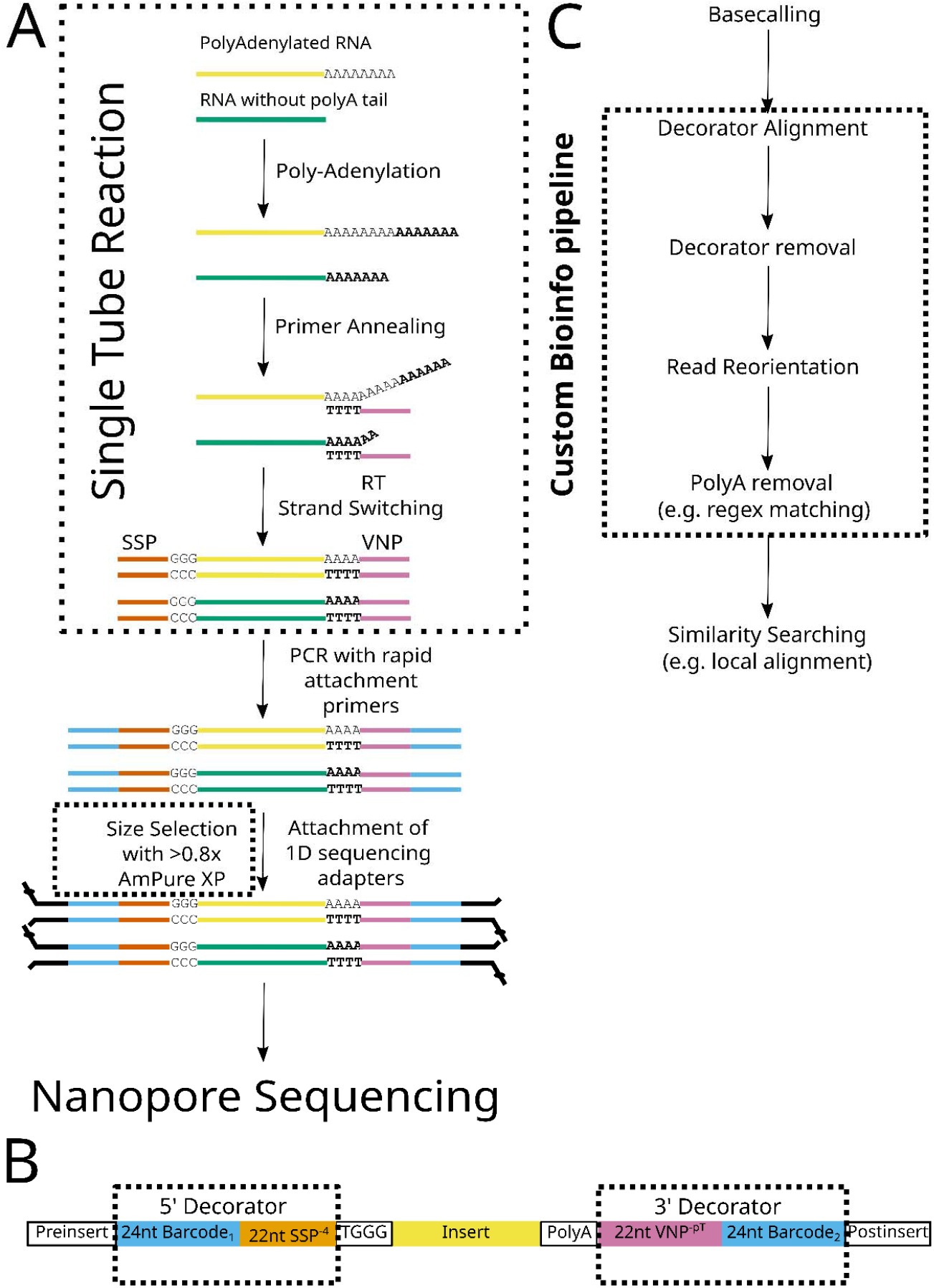
PALS-NS experimental workflow (A), text-based model of a well-formed read (B) and custom bioinformatics pipeline (C). Dashed boxes indicate modifications to biochemical protocols, read models and bioinformatics pipeline.

#### Text model for Nanopore reads

The expected outcome of sequencing is a *well-formed* read containing a single *insert* that is derived from na RNA molecule in the original sample. In such reads (**Figure *1***B) the insert is flanked by the tetrabase TGGG (derived from the 3’ of the SSP) and a poly-A tail at its 5’ and 3’ respectively. External to these features we find the 5’ and 3’ sequence *decorators* (derived from the primers of the PCR during the library preparation) and variable length *pre-insert* and *post-insert* sequences. In the current iteration of the sequences used by ONT in their Smart-seq protocols, the 5’ decorator encompasses a 24nt barcode (Barcode_1_) found in the middle of the reverse PCR primer and the 22 nucleotides of the SSP (sans the tetrabase TGGG, i.e., SSP^−4^). The 3’ decorator is composed of the VNP without its poly-T feature, i.e., VNP^−pT^ and a 24nt Barcode_2_ sequence. At the time of this writing, ONT optionally uses the barcode sequences to multiplex samples (up to 12) for RNAseq; the preinsert and postinsert are derived from the 15nt long sequences flanking these barcodes.

Based on these structural considerations, we thus introduce a text-based model in which the 5’ and 3’ decorator sequences play the role of opening “(“ and closing “)” parentheses in natural language text. During sequencing, motor proteins are attached to both ends of the cDNA molecules in the library, so that molecules may be threaded through the nanopores from either 5’ or 3’ end. If the cDNA molecule is threaded from its 5’ end, it will be sequenced in the 5’→3’ direction and we would read it as (TEXT), but if threaded from its 3’ end it will be sequenced in the 3’→5’ direction we would read it as [rcTEXT]. In these expressions, rc stands for reverse complement, a bracket is the sequence of the decorators when the cDNA is sequenced in the 3’→5’ direction, and TEXT is the sequence of interest comprised of the ACTG alphabet of DNA. The opening bracket “[“ is thus the reverse complement of the closing parenthesis “)”, while the closing bracket “]” is the reverse complement of the opening parenthesis “(“. In our text-model, reads that appear to lack one or both decorators are classified as *partial* and *naked* respectively. It is also possible to *have fusion* reads which contain multiple parentheses and brackets as non-matching pairs e.g. (TEXT]. In such a case, each segment between consecutive parentheses or brackets is classified as a read.

#### Text Based Segmentation of PALS-NS files

We propose a text-based segmentation algorithm (**Figure *1***C) after basecalling the squiggle signal Nanopore files. The algorithm is composed of four steps, which are detailed in the supplement:

1. **Decorator alignment and filtering** using an aligner of choice (in this work we used blastn).
2. **Decorator removal** by extending the decorator alignments to the entire length of the decorator (**Figure *1***B) and insert classification as one of the four *types*: well-formed, fusion, naked, partial. The first round of adapter dimer identification (stage 0 dimers, those whose insert is less than 4 nucleotides) takes place at this stage.
3. **Identification of insert** orientation based on the orientation of the surrounding brackets/parentheses and **re-orientation** of inserts whose sequencing direction has been unambiguously determined by the surrounding decorators, to be in the 5’ →3’ direction.
4. **Removal of poly-A** tails from inserts e.g., via regular expression matching; inserts whose length is smaller than a second user defined threshold, e.g., ten nucleotides are classified as Stage 1 adapter dimers. Longer sequences are used for *blastn*(24) database searches with an e-value threshold of 0.001 to determine similarity.

#### Model for Count Processing

Any given library is hypothesized to generate a set of *mapped* counts *M*_1_, *M*_2_, …, *M_m_* belonging to *m* distinct RNA species, as well as a variable number of nonmapped counts (*M*_0_) and adapter dimers (*M*_−1_). These counts may be modelled as draws from the multinomial distribution:

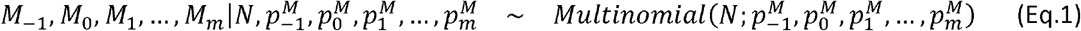

where *N* is the total number of inserts from the library (the library depth), and 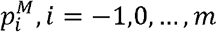 is the fraction of the any given unique RNA species in the library. We can use the properties of the multinomial distribution to analyze:

1. The number of *adapter dimers*, since 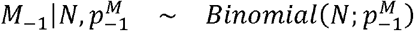
2. The *wasted library depth* (the sum of adapter dimers and non-mapped reads), since

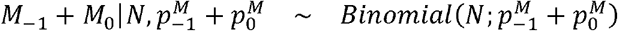
3. The number of *<u>non-mapped reads</u>*, since 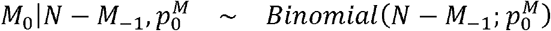
4. The RNA counts of interest *M*_1_, …, *M_n_* by conditioning on the *effective library depth* 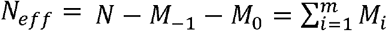 and *the total probability of obtaining a useful read* 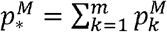 because

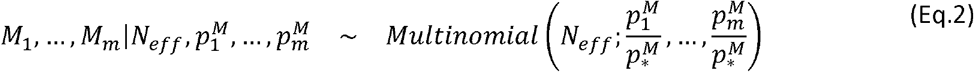
5. The RNA counts of interest 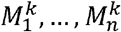 from any *sub-library*, i.e., a subset of the entire library defined by shared characteristics, e.g. the type of insert, and the quality assigned to the corresponding read. This is a straightforward application of (Eq.2) with the counts and probabilities referring to the counts of reads with common features and the effective (sub-)library size is the total count for the particular sub-library.

The probabilities 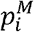 are proportional to the number of cDNA molecules loaded on the flow cell, which is proportional to the number of molecules of each RNA species in the sample (*X_i_*), and the efficiency of the steps of the library preparation. If we assume that the efficiency was the same for all RNAs, then we could simply set 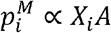, where *A* quantifies the common efficiency of library preparation. Since it is unlikely that this assumption holds true, we are content to write 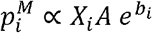, where *b_i_* is a *bias factor* that quantifies the variability in library preparation. In this formulation, the factor *A* yields may be interpreted as a geometric average of the effects of library preparation on the RNA species present in the sample, and the factors *b_i_* as deviations (“random effects”) from this average.

We now introduce a distributional approximation to the model in (Eq.2), that allows to replace the multinomial distribution with the product of independent Poisson random variates (25–28)

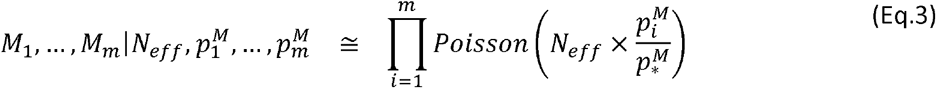

and impose a regression structure on the logarithm of the Poisson mean 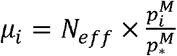:

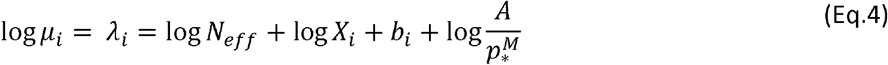

In (Eq.4), the effective library depth, *N_eff_*, is the offset of the regression, and the parameter 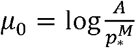 is the overall, *grand* mean. The Poisson models can be extended to account for *overdispersion*, and thus model additional sources of variation that would make RNA counts to be more variable than one would anticipate from Poissonian sampling. The simplest overdispersed model is the Negative Binomial one. In this work, we will be using the Poisson (or the binomial distribution) when the focus is on the performance of the sequencing itself (e.g., analyzing factors affecting the effective library depth), but switching to the Negative Binomial when interest lies in the expression of individual RNAs by aggregating counts over sub-libraries.

##### Exploring bias and dynamic range compression in RNA sequencing via mixed Poisson & Negative Binomial models

An additional complication for the analysis of counts is introduced by the size of the (effective) library depth: while one typically loads a few tens of femtomoles (~10^9^), the sequenced libraries will have a depth ranging between 10^5^ (Flongle) to 10^7^ (MinIon). The impact of the limited depth relative to input is best understood by simulating (Eq.2) for various ranges of *X_i_* for an “average” RNA, i.e. one in which *b_i_* = 0. These simulations shown in **Error! Reference source not found.** illustrate the compression of the dynamic range: RNAs which are present in fraction smaller than the *threshold =Effective Library Depth/ cDNA molecules in Library*) will not generate any counts. The primary means of modelling this effect in a library with known inputs is to replace the logarithm in (Eq.4) by a more general function of the abundance and estimate this function from the data at hand. The simulations of **Error! Reference source not found.** suggest a “stick-breaking” representation, i.e., a linear piecewise function that is constant below the detection threshold and a line with a slope of one for log *X_i_* above the threshold. The modeling task would then be to identify the threshold from counts of RNA species known inputs. However, the presence of noise around the detection threshold suggests that a smoother function (one that “curves”, rather than forming an acute angle around the threshold) would also be a viable option. The regression structure then becomes:

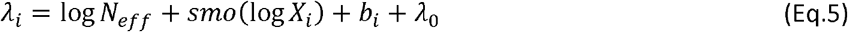

The threshold in this case, will be a “flat” area (a “floor”) over which the counts don’t vary much, if at all, with changes in the input. The function *smo*(·) is given flexibly as a parameterized linear functional (e.g., a cubic or a thin plate spline). The parameters of the spline, which we denote as *θ_s_* and the random effects corresponding to the bias factors may be estimated through penalized regression via Generalized Additive Models(29–31). The latter is a class of modeling tools which allow the data driven estimation of smooth functions and random effects from empirical data. If the input amount (*X_i_*) is known for each RNA, e.g. in the case of synthetic samples of known composition or exogenous spike-ins, then the bias factors *b_i_* could be estimated along with the *smo*(·) from the count data.

### Sources of RNA and Samples

#### Synthetic microRNAs

*We selected 10 microRNAs* (**Error! Reference source not found.**) for the sequencing experiments on the basis of previous work showing their relevance to the author’s research field, kidney biology and pathophysiology(3). MicroRNAs were ordered as single stranded oligos from IDT from the sequences deposited in miRbase(32). One third of the RNAs terminated in ribo-adenines, and half of them included a ribo-adenine within 4 bases of their 3’ end. This was done to test the impact of sequencing errors by the Nanopore device due to the poly-A tails that will be attached to these short RNAs (these analyses will be reported in a future report). Two of the microRNAs were closely related in sequence (200b-5p and 200c-5p) to test the impact of sequencing errors on the identification of microRNAs from the same family. Finally, one sequence (hsa-744-5p) has a ribo-guanine tetraplex; such sequences can form secondary structures which can complicate both synthesis and enzymatic reactions. Additionally, this microRNA has multiple ribo-adenines in its 3’ end, thus providing a special challenge to the proposed workflow. MicroRNAs were aliquoted in stock solutions of 100 μM in TE buffer provided by IDT (10 mM Tris, 0.1 mM EDTA, pH 7.5) and stored in −80oC prior to sequencing. The ten microRNAs randomly allocated to two equimolar pools: a Hi(gh concentration) M(icroRNA) – HiM and a L(ow concentration) M(icroRNA) – LiM one. The final concentration of each RNA in the HiM pool, was double the concentration of each of the microRNAs in the LiM pool (**Error! Reference source not found.).**

#### Synthetic long RNAs

A synthetic spike-mix (ERCC, Thermofisher, Catalog Number 4456740) was used as a source of long RNAs for the sequencing experiments and as a spike in control for the Nanopore experiments involving biological samples. ERCC is a common set of external, unlabeled, polyadenylated RNA controls that was developed by the External RNA Controls Consortium (ERCC)(33) for the purpose of analyzing and controlling for sources of variation in transcriptomic workflows. These transcripts are designed to be 250 to 2,000 nucleotides (nt) in length, which mimic natural eukaryotic mRNAs. The 92 ERCC RNA control transcripts are divided into 4 different subgroups (A-D) of 23 transcripts each. These subgroups are mixed by the vendor to yield a moderate complexity synthetic mix of long transcripts with concentrations that span 6 orders of magnitude (**Error! Reference source not found.).** The length distribution of the ERCC mix and the amount of each RNA vs.. its length is shown in **Error! Reference source not found..** The RNAs in the ERCC and the microRNAs selected, share common subsequences, i.e., half of the length of each short RNAs may be found as “words” inside the longer RNAs as shown in **Error! Reference source not found..** Hence, despite the ERCC being unrelated to biologically derived long RNAs, fragments of the ERCC may be mistaken for short RNAs unless a high stringency sequence database search strategy is utilized. For our experiments we used two separate batches of ERCC (labelled as “A” and “B” in the results tables). The former was maintained in the −80oC to simulate a biological sample under conditions of long-term storage, while the second source was stored as per vendor recommendations in the −20oC and was used for the dilution and the PAP optimization experiments.

#### Construction of synthetic RNA samples (mixes)

The microRNA and ERCC solutions were hand mixed together (**Supplementary Methods**) to generate the following samples:

a. Short RNA sample with a small amount of long, poly-adenylated long RNAs of the ERCC with equimolar mixes of the synthetic miRNAs. In these solutions the microRNAs were presented in a >100fold excess of the ERCC
b. Long RNA samples which contained only the ERCC RNAs
c. A more balanced mix of short and long RNAs in which the short RNAs were present in 5-fold excess over the long RNAs
d. A dilution series 1:1, 1:10, 1:100, 1:1000 of the balanced mix. The preparations at higher dilution were used to simulate low input samples.

These solutions were used to explore the performance of PALS-NS to support a broad range of research agendas: a) poly-A depleted, e.g., short non-coding RNA or tRNA sequencing b) long coding RNA sequencing c) total RNA enriched in poly-adenylated sequences d) ultra-low input samples. RNA from these samples was used as input to the PALS-NS protocol (**Supplementary Methods).**

#### Biological Samples

Total RNA was isolated from the jejunum of ten C57Bl6/J mice from a series of experiments aimed at investigating the impact of different sources and amount of sugar on enteric and kidney physiology. Mice were housed in humidity-, temperature-, and light/dark-controlled rooms and cared for by trained individuals according to the Institutional Animal Care and Use Committees (IACUC) approved-protocols at the University of Cincinnati and the University of New Mexico. Mice were fed either a carbohydrate control or 60% fructose diet (Envigo, Indianapolis, IN) for 5 weeks. Mice were euthanized with an overdose of pentobarbital sodium and jejunum were harvested, cleaned of dietary material, snap frozen and stored at −80o C. Jejunal RNA was extracted using the TRI Reagent method (Molecular Research Center; Cincinnati, OH). RNA samples were stored at −80o C until needed for Illumina sequencing. Two of the isolated samples (one from an animal fed a 60% fructose diet and one fed a carbohydrate control diet) were subjected to Nanopore sequencing using the proposed workflow and the unmodified PCR-cDNA Sequencing Protocol (SQK-PCS109) by Oxford Nanopore Technologies. The biological samples were used to provide an input to the protocol that reflects the composition of naturally occurring RNAs that could be used for library construction.

### Experimental Design

The effects of poly-adenylation in the context of the PALS-NS workflow were analyzed through a series of a 2×2 factorial design experiments utilizing the synthetic RNA samples constructed as detailed previously. In these experiments the two factors considered were: a) addition of short RNA mixes to the ERCC mix vs ERCC mix alone and b) Poly-Adenylation using the protocol in the Supplement vs. Sham Poly-Adenylation. The latter was carried out by incorporating all the elements needed to carry out the poly-adenylation reaction (e.g., reaction buffer, ATP) except the PAP enzyme. In the first 2×2 experiment (“1^st^ 2×2”), the short RNAs were present in >100-fold excess of the ERCC RNA and the entire library was loaded on MinIon flow cells thus saturating the devices. The second of the 2×2 experiments (“2^nd^ 2×2”) was a replicate of the first experiment but only a fixed amount of library was loaded on the flow cells (with the amounts varying by flow cell type). These experiments were used to establish that the PALS-NS protocol can detect short RNAs and paved the way for a focused evaluation of the protocol on the lower capacity (Flongle) flow cells. Subsequently, we evaluated the linear dynamic range of PALS-NS with respect to changes in the molar input of the RNA in a dilution series (“DS”). To construct the DS, we diluted the synthetic mix of short and long RNAs 10-fold, 100-fold and 1000-fold and used these diluted samples as inputs for the library construction. These experiments were controlled by subjecting an ERCC mix to the PALS-NS protocol (only the higher dilution was tested).

### Pre-Sequencing Library Quantitation

#### Synthetic Samples

All synthetic samples were quantitated using High Sensitivity (HS) DNA assays on an Agilent 2100 Bioanalyzer system (Agilent Technologies, Santa Clara, CA). To remain within the assay’s range of quantitation, libraries were diluted either 1:10 or 1:100 with ONT provided Elution Buffer prior to loading the chips. The bioanalyzer output was used to create working libraries of 100 femtomoles (for MinIon flow cells) or 26.12-50 femtomoles (for Flongle flow cells) of cDNA for loading onto the sequencers.

#### Biological Samples

Biological samples were quantitated with a Qubit 3.0 Fluorometer (Life Technologies) using the broad range RNA assay and rudimentary cDNA quality and size information was obtained from an Agilent 2100 Bioanalyzer with a Broad Range DNA Kit (Agilent, USA). For Qubit conversions from ug to picomoles cDNA, the following equation was used:

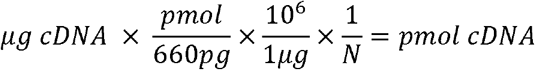

where 660pg is the average molecular weight of a nucleotide pair, and ‘N’ is the predicted number of nucleotides. Upon visual inspection of the Bioanalyzer output, the typical length of the cDNA molecule was 500 bp, giving an estimated input of 200 fmoles to the sequencer.

### Sequencing

#### Nanopore sequencing

Sequencing experiments were done on two Mk1c devices and a single Mk1b device. The criterion for calling a read as low vs.. high quality was a QC score of 8. Fast basecalling (Guppy) was used for all MinIon experiments and high accuracy basecalling for all Flongle experiments. MinIon cells were sequenced for 3 days and Flongle flow cells for 24hrs, but the flow cells were exhausted before then (after approximately 1.5 days for MinIon cells and 9-10 hours for the Flongles). Experiments were run at ONT’s default voltage and temperature settings of −180 mV and 35 degrees Celsius. All flow cells used were of the R9.4.1 chemistry except two flow cells used to sequence biological samples without a polyadenylation step that were of R10.4 chemistry.

#### Illumina Sequencing

The RNA-seq analysis of biological samples was performed by Novogene Bioinformatics Technology Co., Ltd (Beijing, China). Briefly, total RNA isolated from jejunum was subjected to quality control analysis using an Agilent 2100 Bioanalyzer with RNA 6000 Nano Kits (Agilent, USA). After poly A enrichment the samples were fragmented and reverse-transcribed to generate complementary DNA for sequencing. Libraries were sequenced on the HiSeqTM 2500 system (Illumina). Clean reads were aligned to mouse refence genome using Hisat2 v2.0.4.

### Database Mapping

Inserts from synthetic samples were mapped to two different databases of subject sequences: a) ERCC_miRmix, comprised of the 92 ERCC RNAs and the 10 microRNA sequences used to construct the synthetic mixes and b) ERCC_miRBase comprised of the 92 ERCC RNAs and the entire v22.0 mirBase of 48,885 sequences. When analyzing the latter database, we classified reads mapping to the ten microRNAs from different organisms to the human microRNA e.g. mmu-miR-192-5p reads were counted as hsa-miR-192-5p; all other RNAs in miRBase were classified as “NonSpikedRNAs”. To map the biological samples, we created two blast databases a) Mmusculus.39.cDNAncRNA that included all noncoding RNAs and cDNAs from the Genome Reference Consortium Mouse Build 39 and b) Mmusculus.39.cDNAncRNA_spike that enhanced the mouse database with the sequences of the ERCC spike-in mix. The package biomaRt(*34, 35*) was used to map the counts from the biological experiments to ensemble gene ids and eventually gene biotypes.

### Statistical Analysis & Software

A custom bioinformatics pipeline was developed to implement the text-based segmentation algorithm supporting PALS-NS. During segmentation, decorators, inserts and poly-A sequences were individually classified according to the type of the source read (well-formed, partial, fusion, naked) and the quality of the read (“pass” or “fail” as returned by Nanopore’s MinKnow platform). During library mapping, the workflow counted the number of adapter dimers, non-mapped inserts and mapped inserts falling in these eight cross-classifications for each library and generated a text summary with various quality statistics for visual inspection. Result files were from these runs and metadata were loaded into sqlite3 using R’s DBI interface(36). Custom R scripts were written to extract the information from the sqlite3 database for further analyses and deliver pilot implementations of the count processing algorithms, utilizing the GAM modeling package *mgcv*(31) for random effects Poisson and Negative regressions. Insert characteristics were used to fit interaction models in which the effects of experimental factors, polyadenylation vs sham polyadenylation, synthetic RNA source and dilution level were allowed to vary in each by these eight characteristics. These models also allowed us to explore the hypothesis that the representation of distinct RNAs differed in these eight sub-libraries. If the composition of any of these sub-libraries differed from the one found in the gold-standard of the well-formed high-quality reads, then one should strongly consider removing the entire sub-library from further consideration. If on the other hand, the composition does not materially differ, then retaining the sub-library and basing the analyses on the entire set of counts without regard to sub-library type, will not only simplify analyses, but increase the statistical power of experiments based on PALS-NS. Model based cluster analysis with Student-t multivariate components (R package *teigen*(37)) was used to visualize the concordance of libraries generated by different sequencing protocols from the two biological samples.

## Results

### PALS-NS generates inserts of all types with high quality and variable poly-A tails

Experimental conditions for the libraries analyzed in the factorial and dilution experiments are shown in **Error! Reference source not found.**, while the sequencing conditions and overall counts are shown in **Table *1***. We undertook an Analysis of Variance to explore the impact of experimental factors, such as RNA input amount, PAP, flow cell type on the odds of obtaining adapter dimers, non-mapped reads and non-informative reads. The input amount was the most influential factor in these analyses and its effect is shown in **Error! Reference source not found..** All three-quality metrics worsen (positive log-odds ratios) for inputs below ~50 fmoles of RNA input. While adapter dimers continue to decline at higher molar inputs, there are diminishing returns in the other two metrics, largely driven by an increase in the proportion of reads that could not be mapped at the e-value cutoff chosen for these analyses. The vast majority of the experiments in the 2×2, DS as well as the biological samples generated a high number of high quality, well-formed reads (over than half); over 70% of the reads were either partial, or wellformed high quality ones (**Error! Reference source not found.).** A linear regression analysis showed that the average poly-A tail (polyAinter0) from well-formed reads was 15 nucleotides long, while the interrupted poly-A tail types were longer by 5 and 10 nucleotides respectively. The polyAinter0 tails were mostly composed of adenines (98%), as were the polyAinter1 (92%) and polyAinter2 tails (81%). Poly-A tails from fusion and partial reads were shorter by 3 and 2 nucleotides respectively, but their adenine content was lower by 17% and 24% respectively compared to the well-formed reads.

**Table 1.** Sequencing statistics of synthetic libraries created

| Experiment | Library Type | PAP | ERCC Source | Library Replicate | Sequencing Replicate | Dilution | Library Input<sup>†</sup> (fmoles) | Flow Cell | Unique Reads | Unique Inserts | Adapter Dimers N(%) | Noninformative Inserts N(%) | Mapped Inserts N(%) |
| --- | --- | --- | --- | --- | --- | --- | --- | --- | --- | --- | --- | --- | --- |
| 1<sup>st</sup> 2x2 | LiM+HiM+ERCC | + | A | 1 | 1 | 1 | All | FLO-MIN106 | 6575060 | 6660743 | 105908 (1.59) | 3156686 (47.39) | 3504057 (52.61) |
| 2<sup>nd</sup> 2x2 | LiM+HiM+ERCC | + | A | 2 | 1 | 1 | 100.00 | FLO-MIN106 | 2801062 | 2994294 | 6265 (0.21) | 1485668 (49.62) | 1508626 (50.38) |
| 1<sup>st</sup> 2x2 | LiM+HiM+ERCC | - | A | 1 | 1 | 1 | All | FLO-MIN106 | 7116660 | 7160292 | 33470 (0.47) | 251827 (3.52) | 6908465 (96.48) |
| 2<sup>nd</sup> 2x2 | LiM+HiM+ERCC | - | A | 2 | 1 | 1 | 100.00 | FLO-MIN106* | 1044994 | 1051557 | 6831 (0.65) | 75723 (7.20) | 975834 (92.80) |
| 2<sup>nd</sup> 2x2 | LiM+HiM+ERCC | - | A | 2 | 2 | 1 | 35.71 | FLO-FLG001 | 358255 | 367530 | 19566 (5.32) | 51505 (14.01) | 316025 (85.99) |
| 1<sup>st</sup> 2x2 | ERCC | + | A | 1 | 1 | 1 | All | FLO-MIN106 | 9550475 | 9617510 | 142516 (1.48) | 874487 (9.09) | 8743023 (90.91) |
| 2<sup>nd</sup> 2x2 | ERCC | + | A | 2 | 2 | 1 | 34.72 | FLO-FLG001 | 225335 | 245752 | 131694 (53.59) | 245563 (99.92) | 189 (0.08) |
| 1<sup>st</sup> 2x2 | ERCC | - | A | 1 | 1 | 1 | All | FLO-MIN106 | 9272518 | 9325509 | 95731 (1.03) | 664717 (7.13) | 8660792 (92.87) |
| 2<sup>nd</sup> 2x2 | ERCC | - | A | 2 | 1 | 1 | 100.00 | FLO-MIN106* | 1880918 | 1891459 | 14113 (0.75) | 98648 (5.22) | 1792811 (94.78) |
| 2<sup>nd</sup> 2x2 | ERCC | - | A | 2 | 2 | 1 | 34.72 | FLO-FLG001 | 265439 | 267740 | 14895 (5.56) | 37159 (13.88) | 230581 (86.12) |
| DS | ERCC | + | B | 1 | 1 | 1 | 50.00 | FLO-FLG001 | 315945 | 316889 | 1303 (0.41) | 20550 (6.48) | 296339 (93.52) |
| DS | ERCC | + | B | 2 | 1 | 1 | 50.00 | FLO-FLG001 | 198666 | 209200 | 14235 (6.80) | 57692 (27.58) | 151508 (72.42) |
| DS | LiM+HiM+ERCC | + | B | 1 | 1 | 1 | 50.00 | FLO-FLG001 | 17307 | 18839 | 318 (1.69) | 5274 (28.00) | 13565 (72.00) |
| DS | LiM+HiM+ERCC | + | B | 1 | 2 | 1 | 50.00 | FLO-FLG001 | 650850 | 713759 | 18781 (2.63) | 294390 (41.25) | 419369 (58.75) |
| DS | LiM+HiM+ERCC | + | B | 1 | 1 | 10 | 26.12 | FLO-FLG001 | 99903 | 108326 | 19947 (18.41) | 70427 (65.01) | 37899 (34.99) |
| DS | LiM+HiM+ERCC | + | B | 2 | 1 | 10 | 26.12 | FLO-FLG001 | 137583 | 174903 | 67953 (38.85) | 155390 (88.84) | 19513 (11.16) |
| DS | LiM+HiM+ERCC | + | B | 1 | 1 | 100 | 50.00 | FLO-FLG001 | 549899 | 637985 | 352291 (55.22) | 633907 (99.36) | 4078 (0.64) |
| DS | LiM+HiM+ERCC | + | B | 2 | 1 | 100 | 50.00 | FLO-FLG001 | 632184 | 730683 | 357457 (48.92) | 706693 (96.72) | 23990 (3.28) |
| DS | LiM+HiM+ERCC | + | B | 1 | 1 | 1000 | 50.00 | FLO-FLG001 | 189071 | 252980 | 118302 (46.76) | 252341 (99.75) | 639 (0.25) |
| DS | LiM+HiM+ERCC | + | B | 2 | 1 | 1000 | 50.00 | FLO-FLG001 | 236026 | 279255 | 135012(48.35) | 278187 (99.62) | 1068 (0.38) |

DS: Dilution Series, ERCC: External RNA Controls Consortium reference RNA, FLO-MIN106: Minlon Flow Cells, FLO-FLG001 : Flongle Flow Cells, LiM: miRNAs input at lower ratio, HiM: miRNAs input at higher ratio, PAP : Poly A Polymerase, \*reused flow cell , <sup>†</sup>library input estimated via a high sensitivity Bioanalyzer chip (using appropriate dilutions if undiluted library runs failed to yield an estimate of molarity)

### PALS-NS extends Nanopore long read sequencing to short non-coding RNAs

Representation of RNA groups in the 2×2 experiments is shown in **Figure** *3*. ERCC RNAs comprised the bulk (>99%) of all counts a) in the absence of a microRNAs in the sample (sample type ERCC, irrespective of the inclusion of PAP enzyme), and b) when a synthetic mix of microRNAs and ERCC (LiM+HiM+ERCC) was sequenced under sham poly-adenylation (PAP-). The detection rate of microRNAs in the samples which did not include PAP (~0.1% of the effective library depth) is nearly identical to the expected false positive rate (e-value of 0.001) used during the blastn search. When the entire miRBase was used in database searches, the detection rate of the NonSpikedMiRNAs was of the same order of magnitude as that of the other microRNA groups in the sham poly-adenylated samples. The representation of the different RNA groups changed in the LiM+HiM+ERCC samples which were treated with PAP: the ERCC formed the minority of counts as expected from the molarity of the RNA mixes (**Error! Reference source not found.).** Searching against the entire miRBase produced a small number of NonSpikedMiRNAs spurious reads (average 8.2% over all sub-libraries). The counts of the HiM and LiM groups decreased accordingly, suggesting that the spurious reads emanated from sequencing errors that led to the misclassification of short RNAs. Similar results were obtained in the dilution series experiment (**Error! Reference source not found.)** : in the absence of exogenous microRNA input, poly-adenylation does not generate a high rate of false positive short RNA reads, but when such RNAs are included in the RNA source, PALS-NS detects them irrespective of the amount of source RNA.

**Figure 2.**
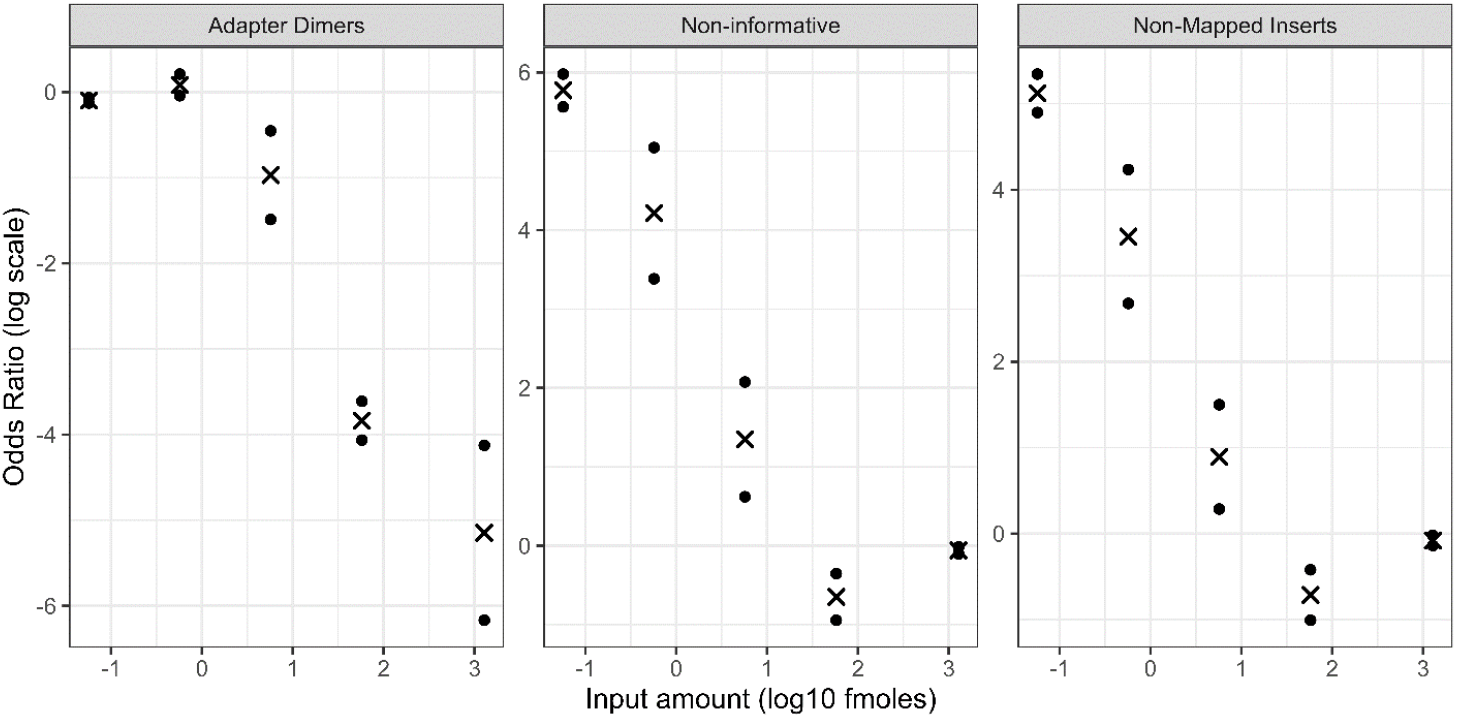
Effect of molar RNA input on quality metrics of the sequencing: Adapter Dimers, Non-informative reads (=adapter dimers and non-mapped reads) and non-mapped inserts (out of the non-adapter dimer inserts). Points : represent individual sequencing run measures, “x” group averages. Plots are given in log-odds scale, with lower (negative) numbers indicating fewer adapter dimers, non-informative and non-mapped inserts.

**Figure 3.**
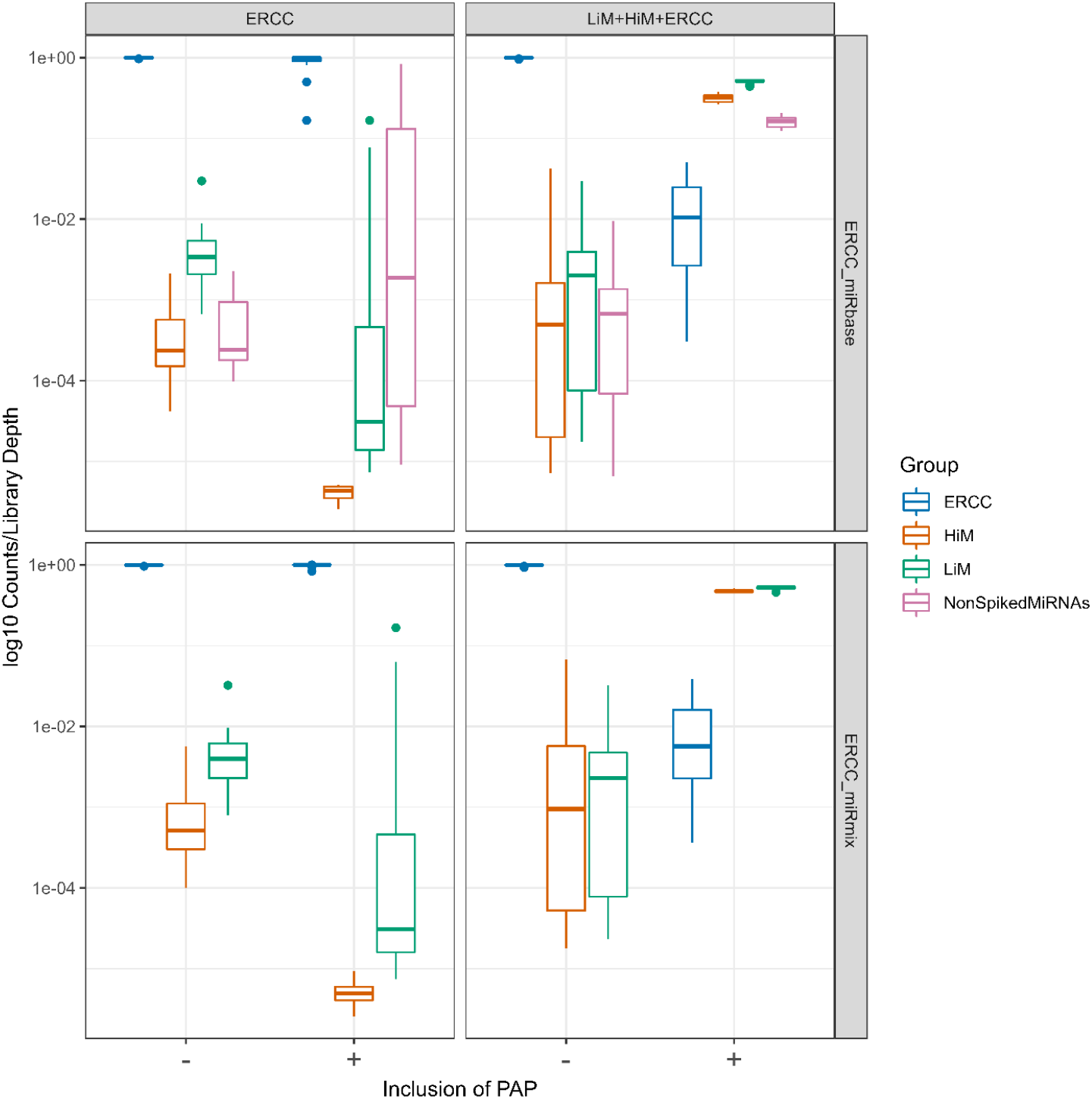
Representation of groups of RNA as a proportion of library depth in the 2×2 experiments; for each sub-library in each sequencing run (a total of eight sub-libraries per library) we calculated the representation of RNAs (counts/effective sub-library depth in log10 scale) according to the group they belonged to: ERCC, HiM or LiM. We cross-classified results according to the RNA input used to construct the library (either ERCC or a mix of LiM, HiM and ERCC) and whether PAP had been included (PAP+) or not (sham poly-adenylation PAP-) after mapping against a sequence library that included the 92 ERCC RNAs and the 10 RNAs used in the mix (ERCC_miRmix). A sensitivity analysis was also performed by mapping against a database of the 92 ERCC and the entire miRbase. In the latter case, counts of RNAs not present in the mix (“NonSpikedMiRNAs”) were tabulated as a separate category.

### PALS-NS segmented inserts can be used to quantify RNA irrespective of insert type and sequencing quality of the source read

Well-formed reads typically accounted for ~55% of all mappable reads, so we tested the hypothesis that the sub-library counts can be grouped together when quantifying RNAs and thus rescue the entire library for quantification. To do so, we fit Poisson regression models to the 2×2 experiment data and included all two way and higher order interactions among the following covariates: sample type (ERCC vs LiM+HiM+ERCC), PAP (sham vs.. actual), read quality (pass vs fail), insert type (one of well-formed, pass, fail, fusion) and RNA group (ERCC, HiM, LiM and when mapping against miRbase NonSpikedMiRNAs). A similar model that included all two way and higher order interactions among dilution, sample type, read quality, insert type, RNA group was also fit to the dilution series. The analysis of variance table for the 2×2 experiments is shown in **Table *2***. The majority of the variance in counts is explained by the RNA Group (i.e. ERCC vs LiM vs HiM), design factors (Sample Type, inclusion of PAP) and interactions between the RNA, Sample Type and PAP. While statistically significant, the interaction terms between insert type, read quality and the experimental factors, explained far less of the variance in counts, and the impact of the latter was quantitatively very small. We illustrate the latter point by generating predictions of the model in **Table *2*** (shown in **Figure** *4*A) for a hypothetical library depth of 10 million inserts. The predicted counts for the RNA groups that were expected to be highly expressed (ERCC in all sham poly-adenylated samples and samples that did not include microRNA input, HiM+LiM in the poly-adenylated LiM+HiM+ERCC samples) were indistinguishable irrespective of the insert type and the quality of the read. On the other hand, predicted counts differed by insert type/read quality for RNA Groups that were either not expected to be present (e.g., the microRNAs in the nonpolyadenylated samples) or anticipated to form only a small fraction of the library (ERCC counts in the LiM+HiM+ERCC libraries that was subjected to polyadenylation). Even in the latter case, the variation in the counts by insert type/read quality was rather small (note how the predicted counts all cluster very closely together in the bottom right panel of **Figure *4***A). Similar results were obtained for the dilution series (**Figure *4***B), indicating that library representation did not materially differ according to insert type and read quality for low input samples.

**Table 2.**
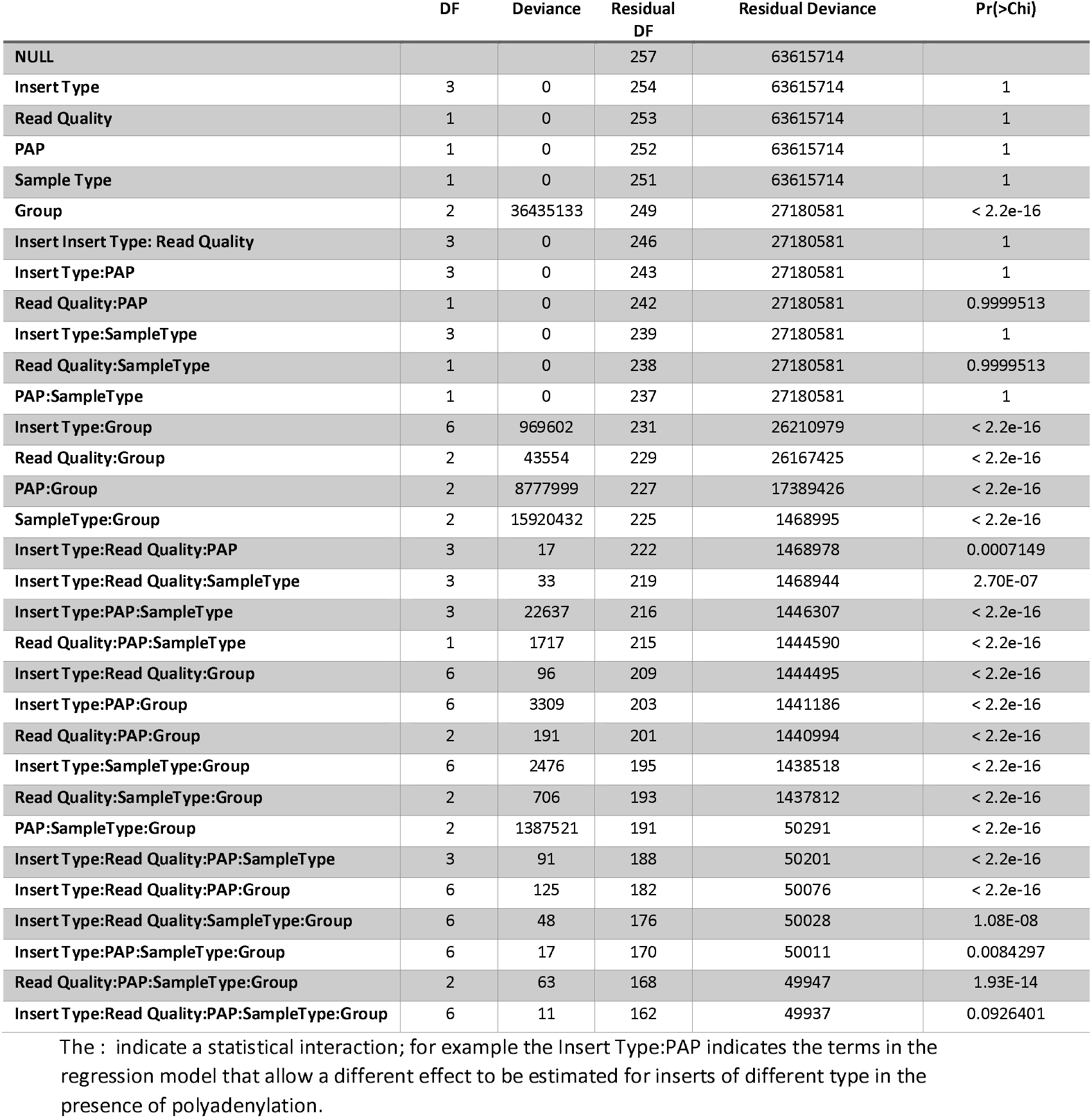
Analysis of variance for the effects of insert type, Read Quality, (RNA) Group and design factors (Polyadenylation, Sample Type, i.e. ERCC vs HiM+LiM+ERCC) in Poisson regressions for the counts from the 2×2 experiments

**Figure 4.**
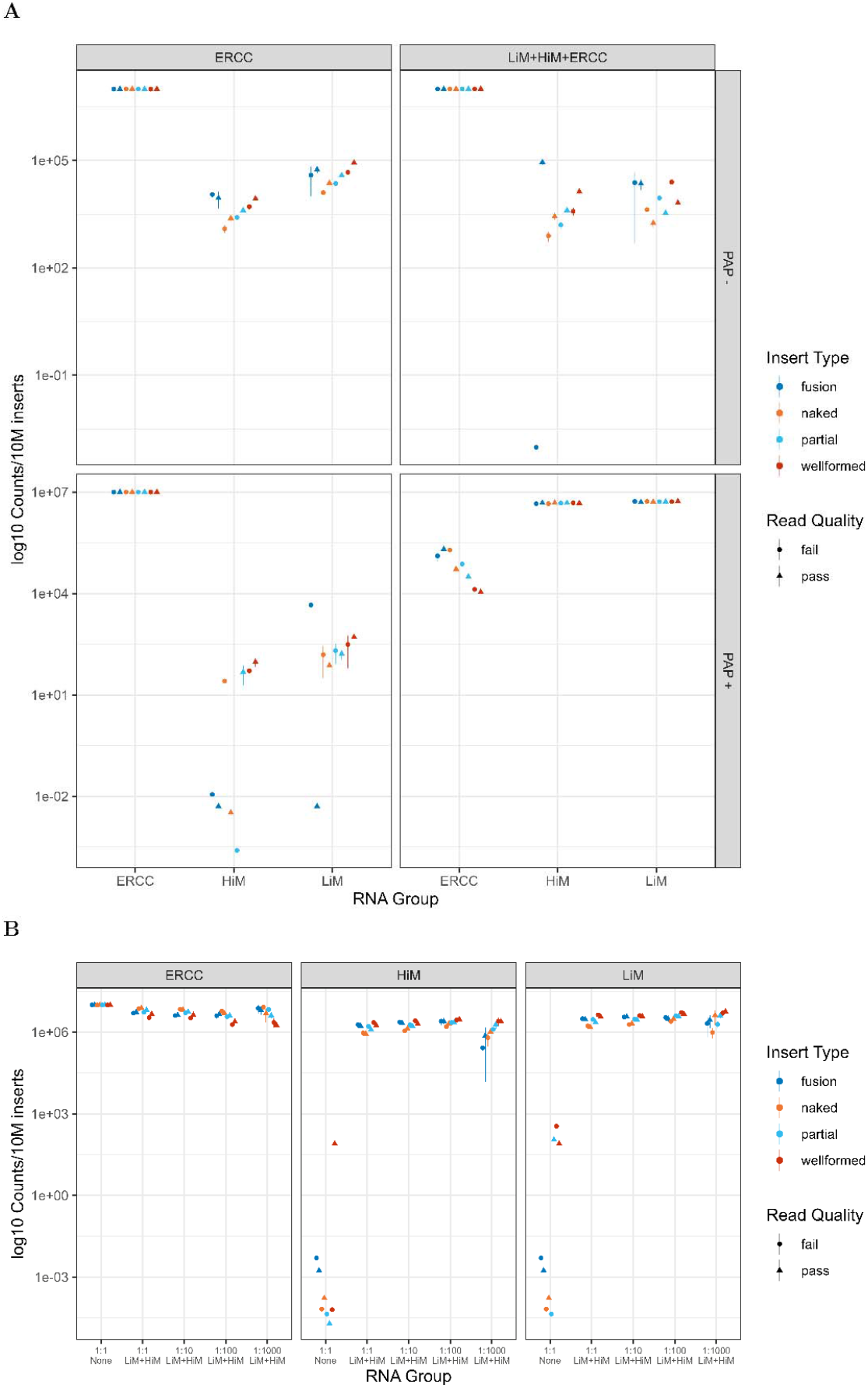
Predicted library representation for a hypothetical depth of 10 million reads by insert type and read quality for the 2×2 experiments (A) and the Dilution Series (B). Sample Types included ERCC (without any microRNA input, “None”), or ERCC with spiked HiM and LiM (LiM+HiM+ERCC in A, LiM+HiM in B)

### PALS-NS quantifies RNAs over eight orders of magnitude of variation in source input while accounting for length and sequence dependence bias

Counts were linearly related to input amount in the absence of PAP (**Figure 5**A), when ERCC was subjected to poly-adenylation (**Figure 5**B) and for the microRNA and ERCC mixture in the 1:1 dilution of the DS experiments (**Figure 5**C). **Figure 5**D-E demonstrate a progressive compression of the dynamic range as the effective library depth (**Table *1***) declined with successive dilutions in the DS. The saturated libraries in the 2×2 experiments (**Figure 5**G) demonstrate a more pronounced form of dynamic compression which was not the result of a decreased library depth (both libraries had > 1.5 million mapped inserts) but was due to the 120-fold excess of microRNAs over ERCCs. In the absence of PAP (**Figure 5**H), there was an inconsistent effect of sequence length on its representation; however there was a small, but definite linear length dependent bias in the presence of PAP (**Figure 5**I): shorter sequences were under-represented relative to their input amount compared to longer sequences. Sequences with length of 20 nucleotides would be underrepresented by a factor of 1.2 log10 ~15.85 times relative to a sequence of 2,020 nucleotides that was present in the same amount as the short sequence in the original RNA sample. The estimated random effects (bias factors) from these analyses are shown in **Error! Reference source not found..** The estimate of the standard deviation of the random effects in log10 scale was 0.482 (95% CI: 0.415 – 0.560), suggesting that 68%, 95%, 99.7% of all RNA counts will be within a factor ranging from 0.329 – 3.03, 0.108 – 9.26, 0.036 – 27.93 respectively relative to the value expected based on their input amount and their length. We also compared the magnitude of the bias factors of the short RNAs in the PAP+ datasets against those obtained in a randomized/degenerate 5’ end 4N ligation based short RNA sequencing protocol(38, 39). To carry out these analyses, we fit the Negative Binomial count model separately to the PAP+ PALS-NS datasets and the publicly available data from the earlier report(38). There was no difference in the bias factors estimated for the common microRNAs (**Error! Reference source not found.**) obtained in these two very different RNAseq protocols (paired t-test for the difference in means p=0.88, Bonett-Seier test of variances for paired samples p=0.67).

**Figure 5.**
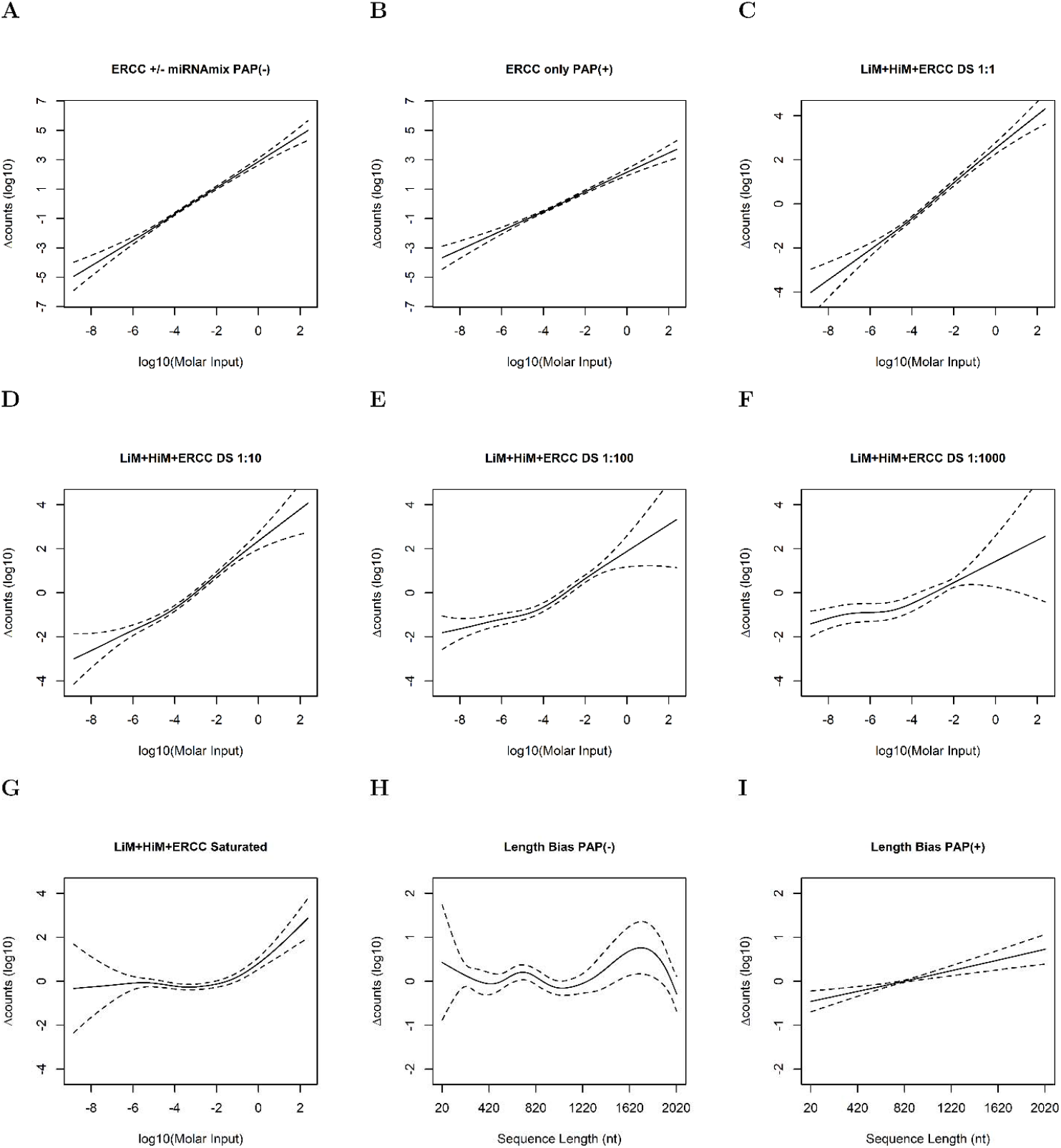
Generalized Additive Model (GAM) Negative Binomial estimates of the variation in sequence count over the 2×2 and dilution series experiment as a function of molar input of each RNA (**A-G**) and sequence length (**H,I**). In additional to the nine functionals, the GAM included a random effect for the (residual) bias factors for each distinct RNA included in these experiments (92 ERCC RNAs and 10 miRNAs for a total of 102 random effects).

### PALS-NS extends the representation of non-coding RNAs in libraries from biological samples

RNA from a control mouse and one fed a high fructose diet were sequenced on an Illumina device, the unmodified SQKPCS109 workflow (denoted as ONT PAP(-) from this point onwards) and PALS-NS yielding libraries with a mapped library depth of 25,722,706/22,686,497 (Illumina), 434,467/717,012 (ONT PAP -) and 4,138,287/8,240,182 (PALS NS respectively when mapping against the Mmusculus.39.cDNAncRNA library). The number of mappable reads for PALS-NS was higher when the Mmusculus.39.cDNAncRNA_spike database was used for searches, i.e. 4,291,187/ 8,525,805 because of the mapping of ERCC reads. The total number of reads obtained on the Nanopore devices were: ~6.7M / 13.8M (Control Diet Sample / High Fructose sample) for the PALS-NS runs and 0.96M/1.3M for the PAP (-) libraries respectively and more than 60% of inserts were mapped. All techniques detected a roughly similar proportion (68-74%) of unique protein coding transcripts and lncRNAs (13-15%) as shown in **Error! Reference source not found..** Other categories of non-coding RNAs (e.g., microRNAs, SnRNAs, ScaRNAs, SnoRNAs) were infrequently detected by Illumina and ONT PAP(-), but rose in frequency in the PALS-NS libraries. While all libraries detected the same protein coding RNAs **Figure 6**A), there was less overlap in the lncRNAs (**Figure 6**B) and much less in the microRNA (**Figure 6**C) and rRNA (**Figure 6**D) categories. Restricting attention to *RNAs that had non-zero counts in at least one library*, correlation was in general strong between libraries obtained by the same method (over 90%, **Error! Reference source not found.).** Correlation was moderate between the Illumina and ONT PAP (-) libraries, and between ONT PAP(-) and PALS-NS (~0.55-0.62) and weak between Illumina and PALS-NS (0.27-0.31). We then explored a) differences in the representation of various RNA species in the three library types and b) dynamic range compression and variably library depth as potential explanations for these variable correlations.

**Figure 6.**
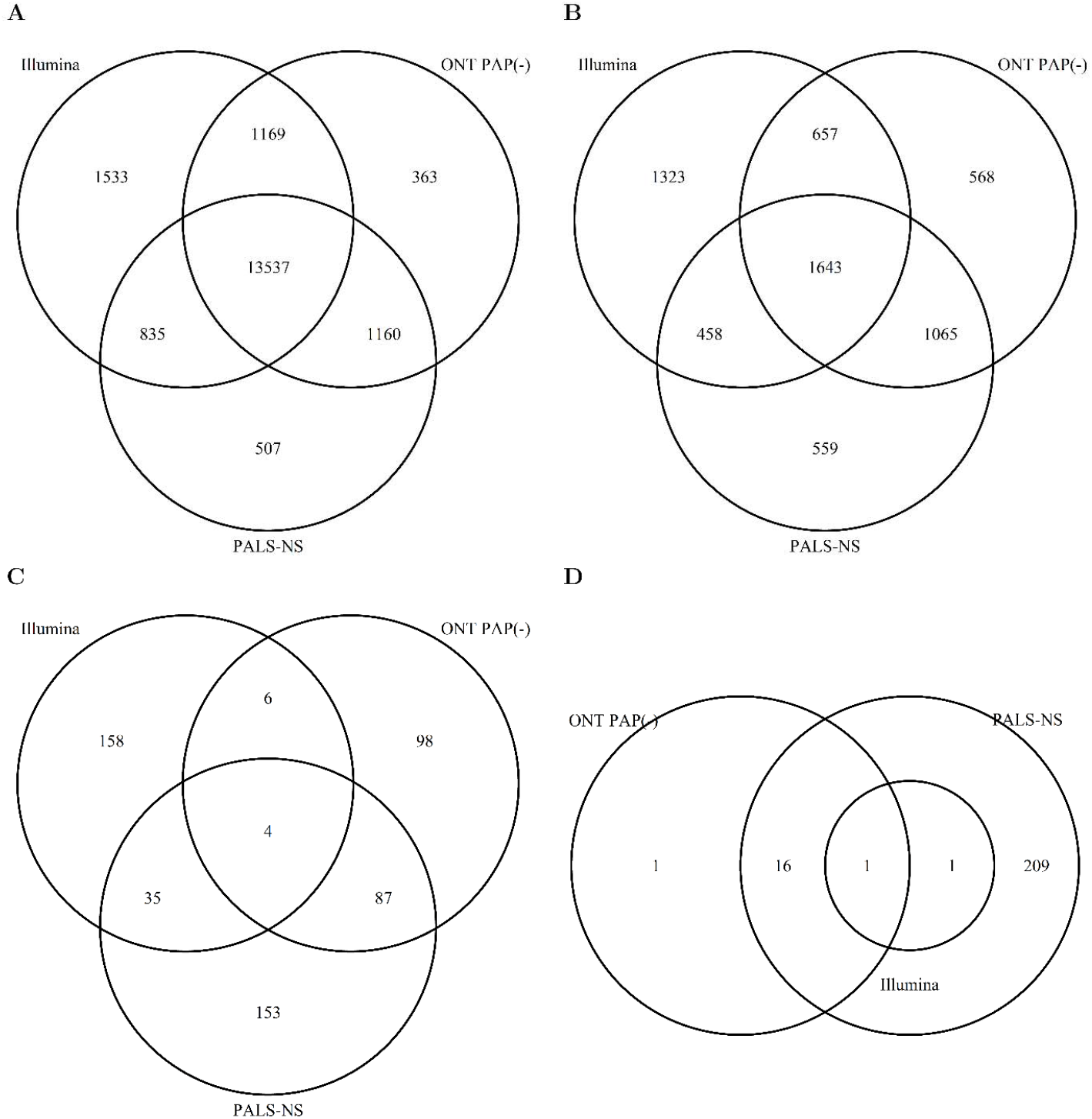
Venn diagram showing the overlap of individual RNAs detected in libraries constructed from a polyA enriched samples (Illumina), long RNA sequencing on a Nanopore device without polyadenylation i.e., ONT PAP(-), and PALS-NS for protein coding RNAs (**A**), long non-coding RNAs (**B**), microRNAs (**C**) and ribosomal RNAs (**D).**

The representation of counts mapping to the Ensembl categories (**Error! Reference source not found.**) revealed some fundamental differences among the Illumina sequencing that used polyA enriched RNA, and the total RNA ONT PAP(-) and PALS-NS libraries. While 98% of the counts from the Illumina libraries mapped to protein coding RNAs, the latter comprised ~84% of the total library depth in the ONT PAP(-) and only 30% of the PALS-NS libraries. Long non-coding RNAs were found in <1% of Illumina libraries, ~3% of ONT PAP(-) but in ~11% of the PALS-NS libraries. Other categories of non-coding RNAs of interest for epigenetics e.g., microRNAs, SnoRNAs, SnRNAs were detected at much higher percentages by PALS-NS. Of note, a significant number of PALS-NS reads mapped to ribosomal RNAs (42%) and mitochondrial transfer RNAs (10%) that were not detected in sizable proportions in the Illumina and ONT PAP (-) sequencing runs. **Table *3*** shows the statistical analysis of the differences in representation of (select) gene biotype categories. Compared to Illumina sequencing, ONT PAP(-) libraries had statistically significant increases in the representation of long non-coding RNAs (lncRNA), microRNAs, mitochondrial RNAs (Mt rRNA and Mt tRNA), ribozymes, small Cajal body RNAs (scaRNA), small nucleolar RNAs (snoRNA), small nuclear (snRNA) and mitochondrial RNAs. Compared to the ONT PAP(-), PALS-NS increased the representation of all non-coding RNAs (except Mt RNA) and decreased as a result the representation of protein coding RNAs.

**Table 3.** Statistical analysis of representation of select gene biotype categories in ONT PAP (-) and PALS-NS libraries versus the same categories in Illumina libraries

| Gene Biotype | ONT PAP (-) vs Illumina |  |  | PALS-NS vs ONT PAP (-) |  |  |
| --- | --- | --- | --- | --- | --- | --- |
|  | RR | 95% CI | p-value | RR | 95% CI | p-value |
| **protein coding** | 0.853 | 0.547 - 1.328 | 0.481 | 0.358 | 0.230 - 0.558 | <0.001 |
| **lncRNA** | 5.203 | 3.340 - 8.107 | <0.001 | 3.519 | 2.259 - 5.482 | <0.001 |
| **miRNA** | 114.617 | 73.337 - 179.133 | <0.001 | 4.360 | 2.796 - 6.798 | <0.001 |
| **misc RNA** | 465.770 | 250.327 - 866.632 | <0.001 | 98.037 | 61.807 - 155.505 | <0.001 |
| **Mt rRNA** | 31.964 | 20.510 - 49.815 | <0.001 | 0.186 | 0.119 - 0.290 | <0.001 |
| **Mt tRNA** | 85.042 | 54.518 - 132.654 | <0.001 | 11.298 | 7.249 - 17.608 | <0.001 |
| **Ribozyme** | 98.336 | 36.730 - 263.268 | <0.001 | 50.425 | 26.121 - 97.342 | <0.001 |
| **rRNA** | 116067.975 | 40619.876 - 331654.747 | <0.001 | 41.778 | 26.808 - 65.108 | <0.001 |
| **scaRNA** | 4.620 | 2.472 - 8.634 | <0.001 | 9.256 | 5.022 - 17.061 | <0.001 |
| **scRNA** | 3252.378 | 99.079 - 106762.804 | <0.001 | 3.240 | 1.754 - 5.983 | <0.001 |
| **snoRNA** | 35.412 | 22.593 - 55.504 | <0.001 | 8.197 | 5.244 - 12.813 | <0.001 |
| **snRNA** | 13.944 | 8.716 - 22.306 | <0.001 | 14.480 | 9.108 - 23.022 | <0.001 |

lncRNA: long non-coding RNA, miRNA: microRNA, Mt rRNA: mitochondrial rRNA, Mt tRNA:
Mitochondrial tRNA, scaRNA: small Cajal body RNA, scRNA: small nuclear RNA, snoRNA: small nucleolar RNA, RR: Relative Ratio, CI : Confidence Interval
Relative Ratios computed on the basis of a Negative Binomial GAM that included all gene biotype categories. Model used a random effect smoother that incorporated gene biotype and sequencing protocol library, as well as random effects at the individual library level.

The effects of dynamic range compression in the biological samples is shown in **Error! Reference source not found..** The floor of detection, the highest count that demarcates the area in which counts don’t vary appreciably by input was visually estimated to be 8 and 16 for the Control Diet and High Fructose libraries. Between the floor and the counts obtained for the most abundant spiked-in RNA, the curve relating molar input to counts appears to undergo two changes in linear slope: the more proximal (to the floor) appears to be at counts of 74 and 155 for the High Fructose and Control Diet libraries, and the terminal one at counts 32 and 74 respectively. To explore the role of the dynamic threshold, we applied a model-based clustering analysis to the counts from non-coding and coding RNAs from the two experiments. The non-coding RNA counts from the 2 samples could be resolved as a cluster of three components (**Figure *7*** A, B). Component A is the cluster of the non-coding RNAs that were not reliably captured by either the Illumina or the PAP (-) ONT library. This cluster appears as a “vertical” ellipse that extends mostly above the floor of PALS-NS for both biological samples, but its projection on the Illumina – PAP (-) plane is oriented along the diagonal because the (low) counts from these two protocols are concordant. Component B is an “horizontal” ellipse of non-coding RNAs with very low counts in the PALS-NS experiments, but with counts that ranged over two orders of magnitude in the Illumina and ONT PAP(-) experiments. These are RNAs whose counts were compressed because of the reduced effective library depth for the complexity of the PALS-NS samples. The correlation of the RNAs mapping to the components A and B is very small as the relevant components are oriented vertically and horizontally respectively. Finally component C includes RNAs that were sequenced above the linear thresholds in both biological samples. The relevant component is oriented along the bottom left – top right direction in the PAP(-) – Illumina and PALS-NS - PAP (-) plane implying a weak positive correlation, but along the top left – bottom right direction in the Illumina - PALS-NS plane implying a negative correlation. Both dynamic range compression, as implied by the component B, and detection of RNAs by PALS-NS that are poorly detectable by the other sequencing protocols (component A) underline the poor correlation among the counts of non-coding RNAs obtainable by PALS-NS, PAP (-) and Illumina. Clustering analysis resolved the coding RNA counts to 4 (Control Diet sample) and 7 (High Fructose sample) components (**Figure *7*** C,D). Coding RNAs captured well by PALS-NS and PAP (-), but not by Illumina map to component A. Components B and D (Control Diet) and B,D and G (High Fructose Diet) are captured by all three sequencing and the relevant components are oriented along a bottom left – top right diagonal indicating a positive and strong correlation. The remaining components (C in the Control Diet, C, E, F in the High Fructose Diet) are RNAs whose counts are highly correlated between the Illumina and PAP (-) libraries, as evidenced by their orientation along the bottom left - top right axis. However, the projection of these components to the 2 PALS-NS planes map at or below the linear thresholds established by the ERCC spike-in analysis, but towards the middle of the Illumina and bottom of the PAP (-) range of counts; these are RNAs whose expression in the PALS-NS libraries was compressed. In summary, the moderate to poor correlation between PALS-NS and either PAP (-) or Illumina is explained by expansion of its repertoire to non-coding RNAs and compression of the dynamic range for the coding RNAs by the over-representation of ribosomal and other non-coding RNAs in the PALS-NS samples.

**Figure 7.**
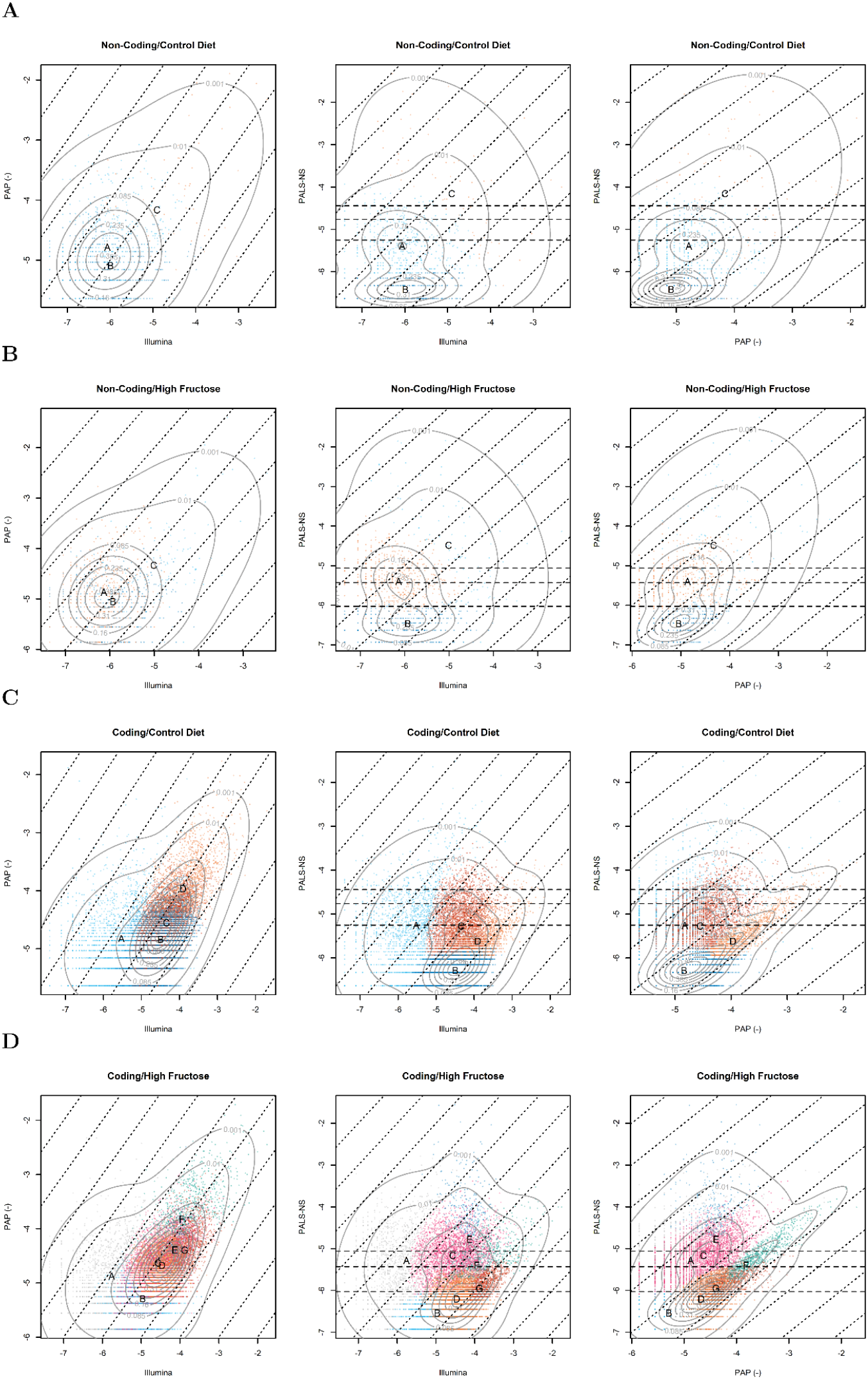
Clustering of counts (expressed as log10 fractions of the library depth for each sequencing) for the two biological samples: Control Diet and High Fructose. To generate these figures, a multivariate clustering algorithm (Teigen) was applied to the three dimensional count data (PALS-NS, PAP (-) and Illumina) of the coding and non-coding RNAs from the two biological samples, for a total of four three dimensional clustering: non-coding RNAs in the Control Diet Sample (**A**), non-coding RNAs in the High Fructose sample (**B**), coding RNAs in the Control Diet Sample (**C**), coding RNAs in the High Fructose sample (D). Each subfigure shows the projection of the density in the three possible planes, the cluster indicator of each count and the centers of the clustering components. The thin dashed oblique lines show the direction of perfect (positive) correlation. The three long dashed horizontal lines are drawn at the linear threshold (upper two) and floor counts of the PALS-NS experiments.

### Length of transcripts sequenced by PALS-NS varies according to the amount of short RNAs present in the sample

In sham-polyadenylated synthetic samples, the length of the ERCC inserts was highly reproducibly and closely tracked the known length of the ERCC irrespective of read quality, or the presence of microRNAs, up to lengths of 784 nucleotides (**Figure 8**A); the length of inserts mapping to longer ERCC RNAs fell below the theoretical length after that point, and only short inserts (below 1000 bases) were recovered for the longest ERCC RNAs. When the synthetic samples were subjected to polyadenylation, the same overall pattern was observed, but there was substantial variability in the insert length within the same length category in the absence of microRNAs, irrespective of the ERCC source and operational PCR parameters (**Figure 8**B, panel ERCC). Inclusion of microRNAs at lower (5x) vs.. higher (>100x) amount relative to the ERCC resulted in longer ERCC mapped inserts irrespective of read quality (**Figure 8**B, panel LiM+HiM+ ERCC). Variation in insert read length did not improve when the synthetic mix of microRNAs and ERCC was diluted (**Figure 8**C); in both the 2×2 experiments (**Figure 8**B) and the DS (**Figure 8**C), the ERCC inserts were shorter than the reference sequence length. The median ERCC insert length in the biological samples appeared to linearly increase in tandem with the reference length up to 784 nucleotides (**Figure 8**D) but declined for longer ERCC sequences. Similarly, the length of inserts mapping to the human transcriptome in the biological samples was longer in the PAP (-) experiments (**Figure *9***A) compared to the PALS-NS runs (**Figure *9***A).

**Figure 8.**
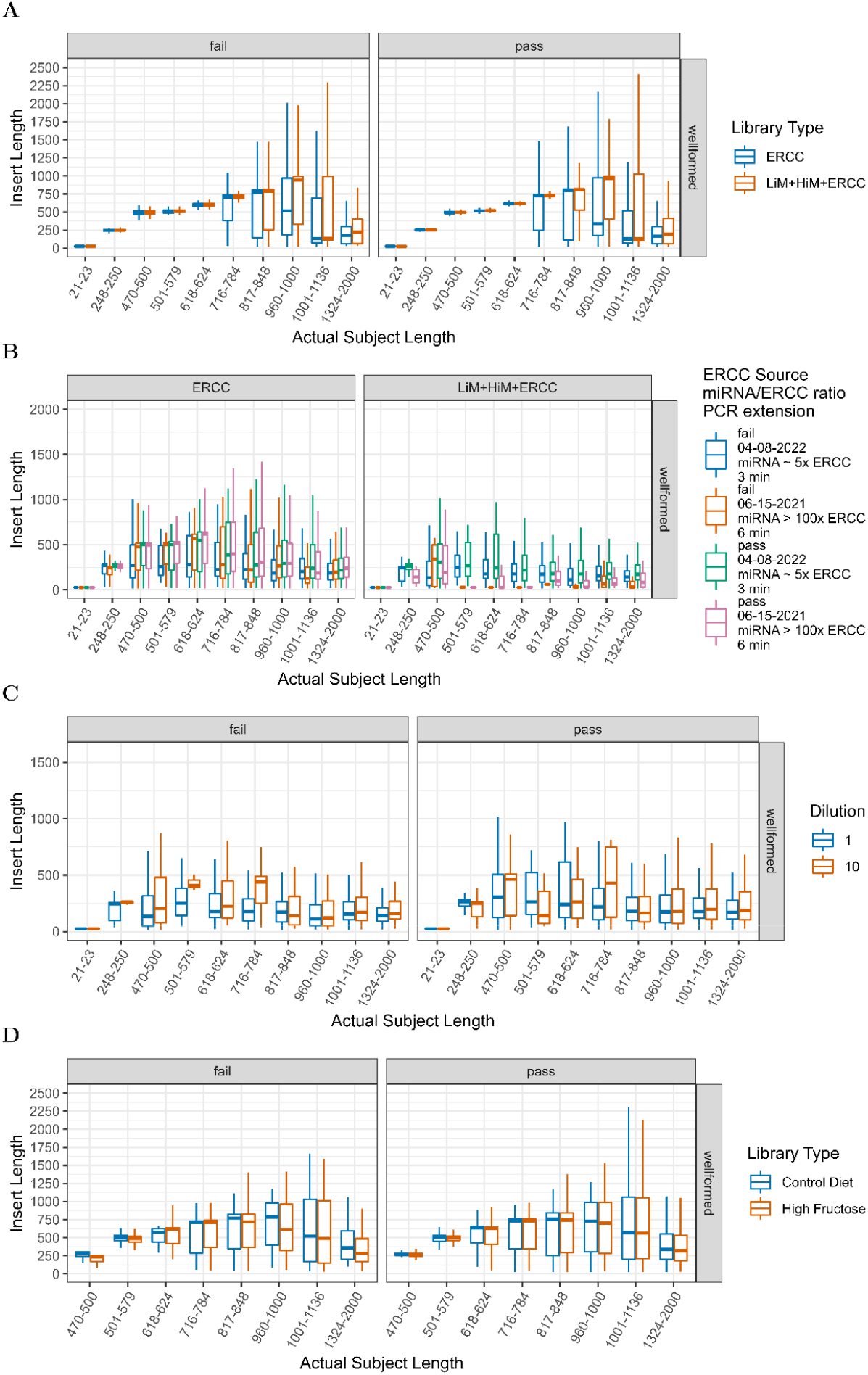
Length of inserts mapping to the ERCC RNAs from the sham poly-adenylated samples (A), the 2×2 and ERCC polyadenylated samples from the DS (B), the LiM+HiM+ERCC samples in the DS (C) and the ERCCs spiked in the two biological samples (D). To generate the graph, ERCC RNAs were grouped together by length, ensuring there at least 4 RNAs per grouping category.

**Figure 9.**
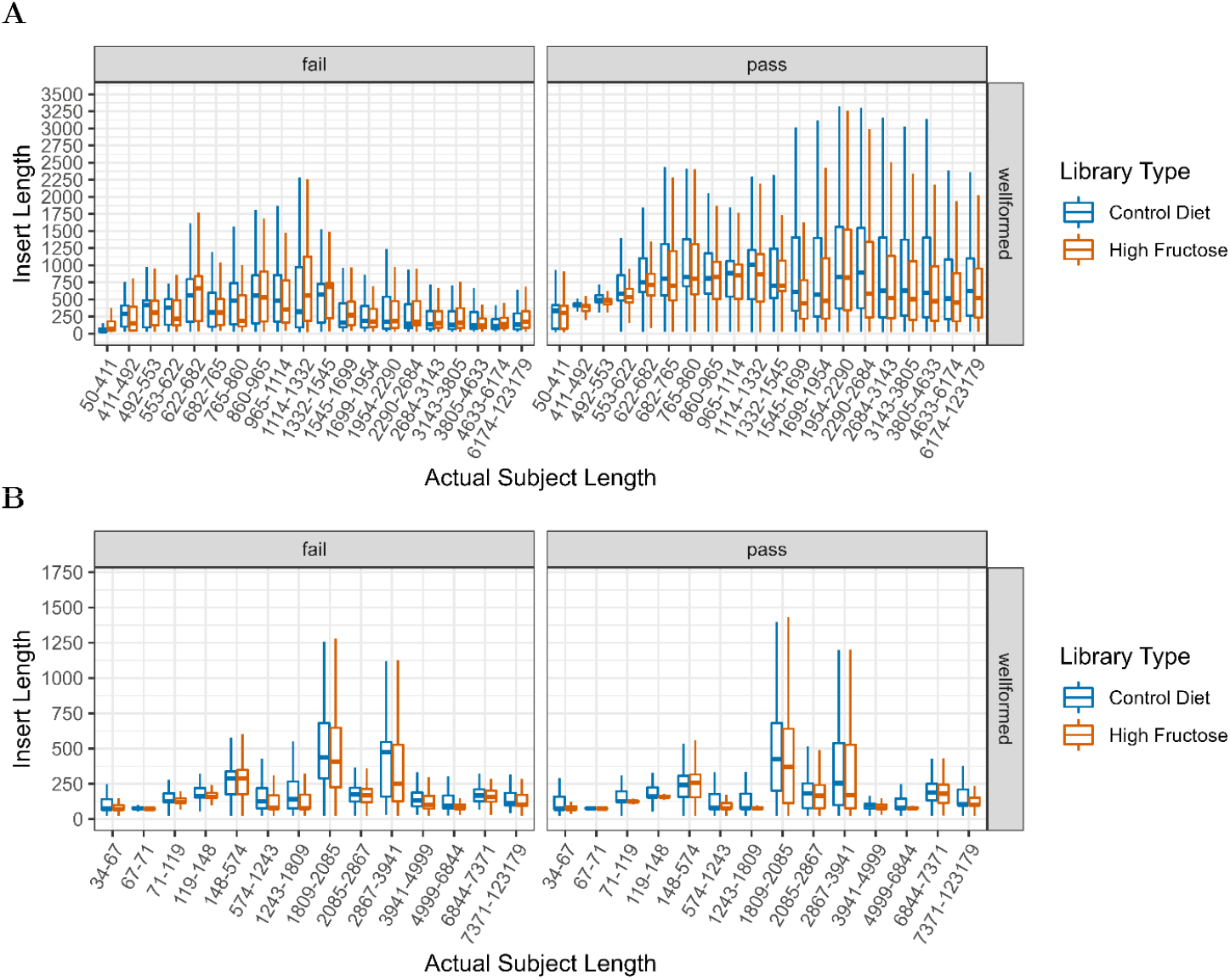
Length of inserts mapping to the human transcriptome in the biological samples from the PAP (-) Nanopore sequencing runs (A) and from the PALS-N protocol (B).

## Discussion

In this work we present a complete solution for the simultaneous profiling of short and long RNAs (either coding and non-coding) from the same library preparation that is sequenced on Nanopore’s device platform. The solution comprises a single tube, single step addition of a homopolymeric tail to the Smart-seq protocol for Nanopore sequencing and a custom bioinformatics pipeline that utilizes a text segmentation model to extract the inserts (RNA sequences in the original sample) from the sequencing reads. Using a variety of synthetic mixes and biological samples we document that the proposed method exhibits exquisite linear dynamic range and will effectively profile both short and long RNAs via Nanopore sequencing. This result, which is due to the innovative combination of biochemistry, a dedicated bioinformatic approach and the single molecule detection capabilities of the Nanopore platform, opens unique opportunities for system biology and biomarker discovery.

### Relation of PALS-NS to previous approaches for short RNA quantification

SMART-seq protocols are ubiquitous since the introduction of the SMART full-length cDNA library construction method more than twenty years ago(21). SMART based protocols utilize a poly-T oligonucleotide to hybridize to the poly-A tail of RNAs, followed by reverse transcription and template switching to synthesize full length first strands. Therefore, RNAs such as microRNAs or transfer RNAs that lack a poly-A tail cannot be analyzed with this technique. Such RNAs can be sequenced via alternative ligation(39–43) and circularization (44, 45) protocols with the former being the default approach to microRNA sequencing. On the other hand, poly-adenylation tagging has been one of the major approaches to quantifying microRNAs by PCR methods(46–51) using universal DNA or Locked Nucleic Acid primers. Our PAP approach is unique by i) clearly separating the PAP and RT reactions in time, but not in space(48), ii) avoiding exposure of polyadenylated RNAs to high temperatures (49, 50, 52) in the presence of magnesium from the PAP reaction buffer that could promote hydrolysis(53) of longer RNAs, iii) moving the entire product to the RT step (51, 54) after cold inactivation and iv) utilizing long, rather than short read RNA sequencing (22, 23). Our approach avoids setting up networks of competitive reaction between the RT and the PAP as both would try to access the 3’ end of the RNAs in the reaction solution. The combination of these technical innovations probably accounts for the high sensitivity of detection of RNAs but also the low sequence and length dependent bias we documented in our analysis. Sequential PAP protocols like ours that execute the PAP and RT steps in a single tube have been reported in the literature previously. However, these protocols though target either qPCR(55) or the Illumina sequencing platforms e.g. CATS/D-Plex (22, 23), Smart-seq-total(19) and thus fail to reap the benefits of long reads, such as reduced length dependent bias and even bias against non-coding RNAs as we discuss below. However, the unique features of Nanopore sequencing requires a text model-based segmentation algorithm to leverage the capabilities of PALS-NS reads which we also discuss below.

### Text model-based segmentation facilitates quantitative analyses of Nanopore Libraries

PALS-NS and other PAP based extensions of SMART-seq protocols (19, 22, 23) for short-read platforms generate a highly structured read, whose “anatomy” was shown in **Figure *1***B. This ideal structure is observed in most, but not all, reads obtained from a single Nanopore experiment as we showed in this work. The variations from the ideal can manifest in various ways, i.e., truncated sequences (naked or partial reads), or fusion/chimeric reads (analogous to pasting one word in the middle of another) and most importantly reversed sequences from cDNAs threaded through the pores in the 3’→5’ direction. If one were to limit attention to reads that conform to the ideal of a well-formed read, one would have to discard on average 45% of the counts in each library, thus further compressing the dynamic range and reducing the quantitative information that can be extracted from Nanopore libraries. To rescue the entire library for quantification, one must accommodate all these variations from the ideal read, which we did by developing a text-based segmentation algorithm that considered all possible deviations from the ideal read.

The first step in our segmentation algorithm is the identification of the location and orientation of the adapters that decorate the insert. The adapter identification step in the PALS-NS operates under similar principles to adapter trimming methods for short-read e.g. cutadapt(56), trimmomatic(57) and long-read sequencing platforms such as Porechop (58), Pychopper (59) and primer-chop(60), i.e. it is a gap alignment based method. The sound statistical properties of the blastn aligner we used control the false positive hits against the decorator sequences (24, 61, 62). Once an alignment has been found, our algorithm extends it to the entire length of the decorators (a form of semi-global alignment). In doing so, we thus fully account for sequencing and basecalling errors that may manifest as internal gaps in the alignment or as abbreviated alignments that don’t cover the entire decorator length. A subtle point in this process, which sets our approach apart from others (59, 60) is our handling of the last four bases in the 3’ end of the 5’ adapter. This is where the RT switches templates and adds non-templated nucleotides. Previous work that examined the template switching junction has shown both sequence dependent bias(63–66) and variable numbers of non-template nucleotides added(64). Therefore, neither the length, nor the composition of the junction can be taken as granted, and thus we opted to retain this feature as part of the insert sequence during text segmentation. This decision guards against the artificial shortening of short RNAs with multiple guanines in their 5’ end, which could have a detrimental impact on their identification during a database similarity search.

The second step in our algorithm, i.e., the reorientation of the insert is an area that has so far attracted limited attention. The only relevant works in this area are ONT’s Pychopper, primer-chop and ReorientExpress(67). The latter is a neural network-based tool that was introduced on the premise that it can identify orientation with higher accuracy than Pychopper and primer-chop. We have not undertaken a direct comparison against these methods, because our text-based segmentation is well suited for the purpose of quantification; only does it rescue all reads (e.g., the current version primer-chop can’t rescue fusion reads), but also appears to do so in a manner that does not compromise quantitation. Hence, the complexity of the dynamic programming algorithm used by pychopper to rescue reads, or the neural network-based method appear to be over-complicated for a task that is solvable by our approach.

The identification and elimination of the polyA tails via regular expression matching is a unique feature of our workflow and should be contrasted to the fixed length poly-A tail used in primer-chop or the fixed length cutadapt methods adopted in CATS/D-Plex and Smart-seq-total. These methods assume that all RNAs will have equal length poly-A, an assumption that is clearly not justified by the variation in the poly-A tails for the RNA species that are naturally poly-adenylated (68, 69), the performance of the PAP enzyme in-vitro(70) or by the non-uniform poly-A tail noted in our own data. On the other hand, our approach allows the poly-A tail to be of variable length, while accommodating a limited number of sequencing errors through the incorporation of non-A patterns in the expressions to be matched. We opine that the meticulous attention to the removal of the decorator and poly-A sequences underline the substantial enhanced mapping rate (over 50% and up to 70%) in the nanogram input libraries, vs.. a figure less than 20% that was previously reported(71). One may wonder whether further enhancements in poly-A detection and removal could improve this mapping rate even further. In that regards, it should be noted that one can use probabilistic methods in the raw, electrical signal (“squiggle”) space to determine the poly-A length (72, 73). While more sophisticated than our proposal, these approaches use specific features of the basecallers in the ONT software platforms and thus will likely have to be calibrated to future implementation of the basecalling suites. On the other hand, as our approach operates on the basecalled sequence, it is independent of the specific basecaller used. However improved approaches to identify and eliminate the poly-A tail, e.g. via different regular expression patterns or Hidden Markov Models (74) are possible, and are currently the subject of investigation by our group.

### PALS-NS extends the scope of long Nanopore reads to short non-coding and long coding and non-coding RNAs

Using synthetic mixes of short (microRNA) and long (ERCC) RNAs we demonstrated that PALS-NS can reliably detect both RNAs in proportion to their input amount. This proportionality is afforded by the non-selectivity of the PAP enzyme for the sequence at the 3’ end of its substrates(75–77). These findings are not likely to be a chance event, because they were obtained in experiments that used sham poly-adenylation and sham short RNA input points to guard against chance variation. The merit of the polyadenylation step in extending the spectrum of Nanopore sequencing was shown in biological samples that were simultaneously sequenced on a short read platform and the unmodified SMART-seq like library preparation kit provided by ONT. PALS-NS detected the entire spectrum of RNAs present in these cellular sources while achieving a balanced ratio between coding and non-coding (e.g., lncRNA/microRNA) counts. An interesting and somewhat novel observation in both the synthetic and biological samples was that the length of the inserts mappable to long RNAs was not always in step with the expected size. Furthermore, the size attained was dependent on both the composition of the sample and the application of a PAP step. These findings were most clearly illustrated in the synthetic samples, and the most plausible explanation is premature template switching in the RT reaction. The mechanism for this premature switching is quite likely a direct competition between short heteroduplexes and partially extended long ones for the RT enzyme. There are several observations that argue in favor of this explanation: firstly this “shortening”, is not observed in the sham poly-adenylated or polyadenylated samples devoid of microRNAs, except for very long RNAs. In the first samples the microRNAs are “invisible” to the RT reaction and in the second samples they are simply not present. In both cases, there are no heteroduplexes to complete for access to the enzyme. Secondly, the amount of shortening depends on the relative ratio of short and long RNAs, demonstrating the quantitative relation expected from a competitive reaction mechanism. Thirdly, this shortening is also observed for pure long RNAs libraries; in that case, intermediate length sequences play the role of short RNAs and limit the length of inserts derived from very long sequences. To our knowledge this is the first time that one reports this finding in RNA Nanopore sequencing, but the mechanism appears general and likely apply to all long-read protocols that utilize a RT step. While the premature termination of RT is certainly an undesirable feature for long read sequencing if the focus is to characterize the *sequence* of the transcripts, it does not impair the ability to *quantitate* RNAs if an alignment method is used to map the inserts against a reference database with statistical rigor. While this competition would seem to limit the case for using a long-read platform, one should note that the amount of shortening will be less in biological samples, than the extreme testing presented by our 2×2 and DS experiments. In fact, using the known composition of the ERCC we observed minimal shortening for RNAs smaller than 624 bases, while more than 50% of inserts deriving from ERCC RNAs between 624-1000bp were of the expected length when the ERCC was spiked in biological samples.

When total RNA is used at the starting material for PALS-NS, the libraries will include numerous counts mapping to “undesirable” RNAs such as ribosomal RNAs and tRNAs; the known discrimination of PAP against 3’ stem loop structures(78–80) was not sufficient to prevent the tailing of molecules with such features despite the presence of multiple other substrates in the biological samples. Hence, if interest lies in the simultaneous profiling of coding and non-coding (lncRNA or microRNAs) the undesirable RNAs should be eliminated to preserve library depth. This can be achieved during sample preparation via RNAse H based treatment or via depletion by CRISPR-Cas9 (19, 81–86) after the cDNA library has been generated. On the other hand, the non-selective nature of poly-adenylation affords the opportunity to develop new sequencing protocols for the Nanopore platform e.g., by combining size selection and poly-A depletion to sequence non-coding RNAs with defined lengths. The ease by which tRNAs are adenylated by this enzyme, a property known since the 1970s, would allow for example the PALS-NS to be used as a novel technique for tRNA quantitative sequencing, an area in dire need of flexible, non-tedious, high-resolution protocols(45). Other possible applications could include sequencing of predominantly non-coding epigenetically relevant RNAs, irrespective of length, after depletion of coding RNAs. The feasibility of such an application was clearly illustrated in the 2×2 experiments that utilized a large excess of microRNAs, to simulate a sample preparation in which the coding RNAs had been depleted prior to library preparation. Considering the emerging role of non-coding short RNAs other than microRNAs(87–90) in the pathogenesis of disease, the PALS-NS protocol offers a unique opportunity to study such RNAs vis-à-vis lncRNAs (6, 7, 91) and short non coding RNAs other than microRNAs. Such studies may allow a better mechanistic understanding(8) of a wide range of disorders and even Nanopore based biomarker assays.

### PALS-NS shows a minimal amount of sequence and length dependent bias for either short or long RNA quantification

Our analyses probed the bias in PALS-NS and of the unmodified cDNA-PCR sequencing workflow for Nanopore devices. This bias may be conceptualized as a deviation of the observed counts of a given cDNA from those expected on the amount of the corresponding RNA present in the original sample. Such deviations may arise from the length of the RNA molecule (“length bias”) or poorly characterized sequence dependent factors (“sequence bias”). In an earlier publication (38), we introduced the term “bias factor” for this deviation and represented it as a random effect in over-dispersed Poisson (negative binomial) in the context of a ligation based, degenerate/randomized end 4N short RNA sequencing protocol. These protocols were the best performing methods in a multi-center evaluation of methods for quantitative microRNA sequencing (39) and performed very well in single cell applications(92). Hence the observation that PALS-NS generates a similar magnitude of bias as one of the highly performing short RNA protocol is encouraging. However, these findings warrants replication and independent verification.

A unique finding of our work relates to the quantification of the length dependent bias of the PALS-NS protocol. This bias, which is a linear, deterministic, function of the *logarithm* of the length of the sequence does not appear to be present in the unmodified RNAseq protocol suggested by ONT, but it does manifest in PALS-NS as an over-representation of longer sequences relative to the shorter ones. We speculate that this bias materializes after poly-adenylation because of the capture of 5’ and mid-sequence fragments of the long RNAs. Lacking a poly-A tail, such fragments would not be represented in a protocol that does not include a poly-adenylation step but inflate the counts of long RNAs during PALS-NS after they acquire such a tail. Stated in other terms, we hypothesize that this is a form of “fragmentation”, length-dependent bias(93–98) similar to that seen in short RNA sequencing platforms. This hypothesis can be resolved by a meticulous alignment analysis of the captured fragments under conditions of sham and actual poly-adenylation. Space considerations preclude us from carrying out such an analysis in this report. Regardless of the mechanisms that lead to such bias, it should be noted that the magnitude is rather small and at least an order of magnitude less than the length dependent bias that is seen with much more expensive sequencing platforms, e.g. the NextSeq and Novaseq(98).

### PALS-NS is a suitable approach for epigenetic research

Our main impetus in developing PALS-NS is to allow simultaneous analysis of non-coding and coding RNAs from a single library preparation for epigenetic research. The molecular biology techniques required to profile short and long RNAs are rather different, thus simultaneous profiling in either bulk(99) or single cell(17) samples requires duplicate workflows, and even different measurement techniques. These may include for example combining RTPCR with microarrays or running separate libraries in the case of sequencing. To our knowledge, the only two publications exploring simultaneous profiling of non-coding and coding RNAs, CATS (22) and Smart-seq-total(19) target the Illumina sequencing platforms. CATS is one of the first papers to explore a PAP protocol and despite the use of older generation RT with substantial RNAseH activity is worth pointing that it achieved a rather large percentage of mappable reads (more than 65), but no information was provided about the microRNAs vs. coding RNAs in the resultant libraries. Like our paper, Smart-seq-total profiled the entire complement of RNA and provides independent verification of the validity of our approach. However, at closer look there are some notable differences between the results reported by the Smart-seq-total investigators and the findings reported here-in that merit consideration. In particular, the ratio of coding/lncRNA/microRNA/snoRNAs/snRNAs as % of the rRNA depleted libraries was reported as 50:1:0.4:1:1 (Figure 1B in (19)), whereas the corresponding ratio was 29:11:2.1:1:0.2 in our data (**Error! Reference source not found.).** While the source of the RNA (bulk in our case, single cells in Smart-seq-total), and sequencing at a different depth with a short read platform may underline these differences, a careful examination of the quantitative aspects of the Smart-seq-total report and this report suggests that such differences may be protocol dependent. Even though the library depths differed between the Smart-seq-total (2.5M reads) and our protocol (4.3M and 8.5M), many reads in our libraries (~40%) mapped to ribosomal RNAs, so that the non-ribosomal library depth we obtained was rather comparable to the Smart-seq-total paper. In support of the latter assertion, examination of the ERCC control counts-molarity curve (Figure S5e in (19)) shows that the dynamic range compression occurred at roughly similar points, i.e. at 3.5 log10 from the least expressed ERCC RNA. Since the number of all RNA biotypes was reduced in the single cell RNA libraries, relative to the bulk, but the library depth was comparable, one would have expected the ratio of read types to be rather similar between Smart-seq-total and PALS-NS. The observation that it is not, suggests that the Smart-seq-total protocol may not be as efficient as PALS-NS in capturing both short and long non-coding RNAs because of biochemistry (tagmentation/pooling, capping reaction), bead clean-up (use of 1.0x volume of beads vs. 1.8x), CRISPR-Cas9 library depletion (19, 85) or the sequencing platform (Novaseq vs Nanopore). While we cannot resolve these differences without further exploration of our protocol in single cell applications, the more balanced representation between coding and non-coding RNAs suggest that the Nanopore based PALS-NS may be more suitable for resolving non-coding RNAs than the Illumina based Smart-seq-total for bulk RNA sequencing.

### PALS-NS demonstrates a very high dynamic range yet requires further optimization for low input samples

A key observation is the extremely high dynamic range of the PALS-NS, which can generate libraries in which the representation of molecules scales linear with abundance over eight orders of magnitude, i.e., much higher than the dynamic range of most, except the very high-end sequencing flow cells. As the relative cost of Nanopore sequencing and the capabilities of the flow cells continue to improve, PALS-NS is well positioned to quantitate RNAs without the library depth limitations of current flow cells. Nevertheless, certain challenges remain to be addressed for low input (3-30 pg) samples, that while easily handled by the protocol, tend to generate a high number of adapter dimers and non-mappable reads. Dimers can be reduced at the magnetic bead clean up stage e.g., by decreasing the ratio of beads to sample volume from 1.8x closer to 1.0x at the expense of losing a variable amount of the short RNA derived inserts. On the other hand, the high percentage of non-mappable reads may require optimization of the PAP and RT steps as these reads likely originate at the interface of these reactions. Previous work has shown that while the mapping rate of Maxima Minus H derived libraries will be in the 85-90% range(65) for ng input (also observed in our work), the mapping rate will decline to ~50% in the pigogram range. Mapping of low input PALS-NS libraries such as the 1:100 and 1:1000 diluted synthetic mixes was much lower, suggesting that the high number of non-mappable reads may originate at this stage. It should be noted that similar observations, i.e. high dimers and a large proportion of nonmappable reads were made in an evaluation of the CATS protocol for the ultimate low input sample, i.e. single cell microRNA sequencing(92). Hence, extending the scope of PALS-NS to low input or even single cell RNA sequencing applications will require further kinetic(19) or input(22, 23) optimization of the PAP reaction and possibly exploration of alternative reverse transcriptase enzymes.

## Conclusions

1. PALS-NS is capable of simultaneously profiling short and long RNAs from a single tube reaction through a simple PAP modification of existing SMART-seq protocols and associated bioinformatics workflow using Nanopore sequences
2. PALS-NS extends the dynamic range of reads detection to non-coding RNAs with limited length and sequence-dependent bias.
3. Bias for short RNAs is comparable to the gold standard Illumina protocol (4N) developed by NIH’s exRNA consortium
4. The entire complement of RNAs in biological samples is profiled by PALS-NS in bulk RNAseq applications
5. Future adaptations of the protocol may extend its scope for (ultra-)low input samples and single cell RNA sequencing.

## Supporting information

Supplemental Methods and Results

## Acknowledgements and Financial Support

This grant was financially supported by DCI Inc. #C-3765 “A community based study of the Epidemiology of CKD in rural New Mexico” to CA. Research performed at the University of New Mexico Clinical and Translational Science Center (CTSC) was supported by an award from the National Center for Advancing Translational Sciences, National Institutes of Health under grant number UL1TR001449.

## References

1. Ben-Dov, I.Z., Tan, Y.-C., Morozov, P., Wilson, P.D., Rennert, H., Blumenfeld, J.D. and Tuschl, T. (2014) Urine microRNA as potential biomarkers of autosomal dominant polycystic kidney disease progression: description of miRNA profiles at baseline. PLoS ONE, 9, e86856.

2. Ben-Dov, I.Z., Whalen, V.M., Goilav, B., Max, K.E.A. and Tuschl, T. (2016) Cell and Microvesicle Urine microRNA Deep Sequencing Profiles from Healthy Individuals: Observations with Potential Impact on Biomarker Studies. PLOS ONE, 11, e0147249.

3. Shaffi, S.K., Galas, D., Etheridge, A. and Argyropoulos, C. (2018) Role of MicroRNAs in Renal Parenchymal Diseases—A New Dimension. International Journal of Molecular Sciences, 19, 1797.

4. Nassirpour, R., Raj, D., Townsend, R. and Argyropoulos, C. (2016) MicroRNA biomarkers in clinical renal disease: from diabetic nephropathy renal transplantation and beyond. Food Chem. Toxicol., 10.1016/j.fct.2016.02.018.

5. Liu, C., Ma, K., Zhang, Y., He, X., Song, L., Chi, M., Han, Z., Li, G., Zhang, Q. and Liu, C. (2022) Kidney diseases and long non-coding RNAs in the limelight. Frontiers in Physiology, 13.

6. Lekka, E. and Hall, J. (2018) Noncoding RNAs in disease. FEBS Lett, 592, 2884–2900.

7. Statello, L., Guo, C.-J., Chen, L.-L. and Huarte, M. (2021) Gene regulation by long non-coding RNAs and its biological functions. Nat Rev Mol Cell Biol, 22, 96–118.

8. Fabbri, M., Girnita, L., Varani, G. and Calin, G.A. (2019) Decrypting noncoding RNA interactions, structures, and functional networks. Genome Res., 29, 1377–1388.

9. Wang, S., Xia, P., Zhang, L., Yu, L., Liu, H., Meng, Q., Liu, S., Li, J., Song, Q., Wu, J., et al. (2019) Systematical Identification of Breast Cancer-Related Circular RNA Modules for Deciphering circRNA Functions Based on the Non-Negative Matrix Factorization Algorithm. Int J Mol Sci, 20, 919.

10. Huang, Y.-A., Chan, K.C.C. and You, Z.-H. (2018) Constructing prediction models from expression profiles for large scale lncRNA–miRNA interaction profiling. Bioinformatics, 34, 812–819.

11. Guo, D., Fan, Y., Yue, J.-R. and Lin, T. (2021) A regulatory miRNA–mRNA network is associated with transplantation response in acute kidney injury. Human Genomics, 15, 69.

12. Zhang, Y., Zhang, L., Wang, Y., Ding, H., Xue, S., Qi, H. and Li, P. (2019) MicroRNAs or Long Noncoding RNAs in Diagnosis and Prognosis of Coronary Artery Disease. Aging Dis, 10, 353–366.

13. de Gonzalo-Calvo, D., Vea, A., Bär, C., Fiedler, J., Couch, L.S., Brotons, C., Llorente-Cortes, V. and Thum, T. (2019) Circulating non-coding RNAs in biomarker-guided cardiovascular therapy: a novel tool for personalized medicine? European Heart Journal, 40, 1643–1650.

14. Yang, Y., Li, Y., Yang, H., Guo, J. and Li, N. (2021) Circulating MicroRNAs and Long Non-coding RNAs as Potential Diagnostic Biomarkers for Parkinson’s Disease. Front Mol Neurosci, 14, 631553.

15. Yang, L., Wang, B., Ma, L. and Fu, P. (2022) An Update of Long-Noncoding RNAs in Acute Kidney Injury. Frontiers in Physiology, 13.

16. Volovat, S.R., Volovat, C., Hordila, I., Hordila, D.-A., Mirestean, C.C., Miron, O.T., Lungulescu, C., Scripcariu, D.V., Stolniceanu, C.R., Konsoulova-Kirova, A.A., et al. (2020) MiRNA and LncRNA as Potential Biomarkers in Triple-Negative Breast Cancer: A Review. Frontiers in Oncology, 10.

17. Wang, N., Zheng, J., Chen, Z., Liu, Y., Dura, B., Kwak, M., Xavier-Ferrucio, J., Lu, Y.-C., Zhang, M., Roden, C., et al. (2019) Single-cell microRNA-mRNA co-sequencing reveals non-genetic heterogeneity and mechanisms of microRNA regulation. Nat Commun, 10, 95.

18. Xiao, Z., Cheng, G., Jiao, Y., Pan, C., Li, R., Jia, D., Zhu, J., Wu, C., Zheng, M. and Jia, J. (2018) Holo-Seq: single-cell sequencing of holo-transcriptome. Genome Biology, 19, 163.

19. Isakova, A., Neff, N. and Quake, S.R. (2021) Single-cell quantification of a broad RNA spectrum reveals unique noncoding patterns associated with cell types and states. Proceedings of the National Academy of Sciences, 118, e2113568118.

20. Wang, Y., Zhao, Y., Bollas, A., Wang, Y. and Au, K.F. (2021) Nanopore sequencing technology, bioinformatics and applications. Nat Biotechnol, 39, 1348–1365.

21. Zhu, Y. y., Machleder, E. m., Chenchik, A., Li, R. and Siebert, P. d. (2001) Reverse Transcriptase Template Switching: A SMART™ Approach for Full-Length cDNA Library Construction. BioTechniques, 30, 892–897.

22. Turchinovich, A., Surowy, H., Serva, A., Zapatka, M., Lichter, P. and Burwinkel, B. (2014) Capture and Amplification by Tailing and Switching (CATS). RNA Biol, 11, 817–828.

23. D-Plex Small RNA-seq Kit. Small RNA library preparation kit for Illumina® sequencing (2021).

24. Altschul, S.F., Gish, W., Miller, W., Myers, E.W. and Lipman, D.J. (1990) Basic local alignment search tool. J Mol Biol, 215, 403–410.

25. Hodges, J.L. and Le Cam, L. (1960) The Poisson Approximation to the Poisson Binomial Distribution. The Annals of Mathematical Statistics, 31, 737–740.

26. McDonald, D.R. (1980) On the Poisson Approximation to the Multinomial Distribution. The Canadian Journal of Statistics / La Revue Canadienne de Statistique, 8, 115–118.

27. Blanc, A.S. (1991) On the Poisson approximation for some multinomial distributions. Statistics & Probability Letters, 11, 1–6.

28. Poor, H.V. (1991) The maximum difference between the binomial and Poisson distributions. Statistics & Probability Letters, 11, 103–106.

29. Wood, S.N. (2011) Fast stable restricted maximum likelihood and marginal likelihood estimation of semiparametric generalized linear models. Journal of the Royal Statistical Society: Series B (Statistical Methodology), 73, 3–36.

30. Wood, S.N., Goude, Y. and Shaw, S. (2015) Generalized additive models for large data sets. J. R. Stat. Soc. C, 64, 139–155.

31. Wood, S.N. (2017) Generalized Additive Models: An Introduction with R, Second Edition 2nd edition. Chapman and Hall/CRC, Boca Raton.

32. Kozomara, A. and Griffiths-Jones, S. (2014) miRBase: annotating high confidence microRNAs using deep sequencing data. Nucl. Acids Res., 42, D68–D73.

33. External RNA Controls Consortium (2005) Proposed methods for testing and selecting the ERCC external RNA controls. BMC Genomics, 6, 150.

34. Zhang, J., Haider, S., Baran, J., Cros, A., Guberman, J.M., Hsu, J., Liang, Y., Yao, L. and Kasprzyk, A. (2011) BioMart: a data federation framework for large collaborative projects. Database, 2011, bar038–bar038.

35. Durinck, S., Moreau, Y., Kasprzyk, A., Davis, S., De Moor, B., Brazma, A. and Huber, W. (2005) BioMart and Bioconductor: a powerful link between biological databases and microarray data analysis. Bioinformatics, 21, 3439–3440.

36. R Special Interest Group on Databases (R-SIG-DB), Wickham, H. and Müller, K. (2022) DBI: R Database Interface.

37. Andrews, J.L., Wickins, J.R., Boers, N.M. and McNicholas, P.D. (2018) teigen: An R Package for Model-Based Clustering and Classification via the Multivariate t Distribution. Journal of Statistical Software, 83, 1–32.

38. Argyropoulos, C., Etheridge, A., Sakhanenko, N. and Galas, D. (2017) Modeling bias and variation in the stochastic processes of small RNA sequencing. Nucleic Acids Research, 45, e104.

39. Giraldez, M.D., Spengler, R.M., Etheridge, A., Godoy, P.M., Barczak, A.J., Srinivasan, S., De Hoff, P.L., Tanriverdi, K., Courtright, A., Lu, S., et al. (2018) Comprehensive multi-center assessment of small RNA-seq methods for quantitative miRNA profiling. Nat Biotechnol, 36, 746–757.

40. Baran-Gale, J., Kurtz, C.L., Erdos, M.R., Sison, C., Young, A., Fannin, E.E., Chines, P.S. and Sethupathy, P. (2015) Addressing Bias in Small RNA Library Preparation for Sequencing: A New Protocol Recovers MicroRNAs that Evade Capture by Current Methods. Front. Genet., 10.3389/fgene.2015.00352.

41. Dard-Dascot, C., Naquin, D., d’Aubenton-Carafa, Y., Alix, K., Thermes, C. and van Dijk, E. (2018) Systematic comparison of small RNA library preparation protocols for next-generation sequencing. BMC Genomics, 19, 118.

42. Wong, R.K.Y., MacMahon, M., Woodside, J.V. and Simpson, D.A. (2019) A comparison of RNA extraction and sequencing protocols for detection of small RNAs in plasma. BMC Genomics, 20, 446.

43. van Dijk, E.L. and Thermes, C. (2021) A Small RNA-Seq Protocol with Less Bias and Improved Capture of 2Ͱ-O-Methyl RNAs. In McMahon, M. (ed), RNA Modifications: Methods and Protocols, Methods in Molecular Biology. Springer US, New York, NY, pp. 153–167.

44. Barberán-Soler, S., Vo, J.M., Hogans, R.E., Dallas, A., Johnston, B.H. and Kazakov, S.A. (2018) Decreasing miRNA sequencing bias using a single adapter and circularization approach. Genome Biology, 19, 105.

45. Behrens, A., Rodschinka, G. and Nedialkova, D.D. (2021) High-resolution quantitative profiling of tRNA abundance and modification status in eukaryotes by mim-tRNAseq. Molecular Cell, 81, 1802–1815.e7.

46. Jensen, S.G., Lamy, P., Rasmussen, M.H., Ostenfeld, M.S., Dyrskjøt, L., Ørntoft, T.F. and Andersen, C.L. (2011) Evaluation of two commercial global miRNA expression profiling platforms for detection of less abundant miRNAs. BMC Genomics, 12, 435.

47. Forero, D.A., González-Giraldo, Y., Castro-Vega, L.J. and Barreto, G.E. (2019) qPCR-based methods for expression analysis of miRNAs. BioTechniques, 67, 192–199.

48. Shi, R. and Chiang, V.L. (2005) Facile means for quantifying microRNA expression by real-time PCR. BioTechniques, 39, 519–525.

49. Balcells, I., Cirera, S. and Busk, P.K. (2011) Specific and sensitive quantitative RT-PCR of miRNAs with DNA primers. BMC Biotechnology, 11, 70.

50. Busk, P.K. (2015) Method for quantification of small RNA species.

51. Kang, K., Zhang, X., Liu, H., Wang, Z., Zhong, J., Huang, Z., Peng, X., Zeng, Y., Wang, Y., Yang, Y., et al. (2012) A Novel Real-Time PCR Assay of microRNAs Using S-Poly(T), a Specific Oligo(dT) Reverse Transcription Primer with Excellent Sensitivity and Specificity. PLOS ONE, 7, e48536.

52. Busk, P. (2010) Method for Quantification of Small Rna Species.

53. Tenhunen, J. (1989) Hydrolysis of single-stranded RNA in aqueous solutions—effect on quantitative hybridizations. Molecular and Cellular Probes, 3, 391–396.

54. Niu, Y., Zhang, L., Qiu, H., Wu, Y., Wang, Z., Zai, Y., Liu, L., Qu, J., Kang, K. and Gou, D. (2015) An improved method for detecting circulating microRNAs with S-Poly(T) Plus real-time PCR. Sci Rep, 5, 15100.

55. Lee, Y.-H., Hsueh, Y.-W., Peng, Y.-H., Chang, K.-C., Tsai, K.-J., Sun, H.S., Su, I.-J. and Chiang, P.-M. (2017) Low-cell-number, single-tube amplification (STA) of total RNA revealed transcriptome changes from pluripotency to endothelium. BMC Biology, 15, 22.

56. Martin, M. (2011) Cutadapt removes adapter sequences from high-throughput sequencing reads. EMBnet.journal, 17, 10–12.

57. Bolger, A.M., Lohse, M. and Usadel, B. (2014) Trimmomatic: a flexible trimmer for Illumina sequence data. Bioinformatics, 30, 2114–2120.

58. Wick, R. (2018) Porechop.

59. Pychopper (2022).

60. Frith, M. (2022) primer-chop.

61. Mitrophanov, A.Yu. and Borodovsky, M. (2006) Statistical significance in biological sequence analysis. Briefings in Bioinformatics, 7, 2–24.

62. Pagni, M. and Jongeneel, C.V. (2001) Making sense of score statistics for sequence alignments. Briefings in Bioinformatics, 2, 51–67.

63. Zajac, P., Islam, S., Hochgerner, H., Lönnerberg, P. and Linnarsson, S. (2013) Base Preferences in Non-Templated Nucleotide Incorporation by MMLV-Derived Reverse Transcriptases. PLOS ONE, 8, e85270.

64. Wulf, M.G., Maguire, S., Humbert, P., Dai, N., Bei, Y., Nichols, N.M., Corrêa, I.R. and Guan, S. (2019) Non-templated addition and template switching by Moloney murine leukemia virus (MMLV)-based reverse transcriptases co-occur and compete with each other. Journal of Biological Chemistry, 294, 18220–18231.

65. Jia, E., Shi, H., Wang, Y., Zhou, Y., Liu, Z., Pan, M., Bai, Y., Zhao, X. and Ge, Q. (2021) Optimization of library preparation based on SMART for ultralow RNA-seq in mice brain tissues. BMC Genomics, 22, 809.

66. Wulf, M.G., Maguire, S., Dai, N., Blondel, A., Posfai, D., Krishnan, K., Sun, Z., Guan, S. and Corrêa, I.R. (2022) Chemical capping improves template switching and enhances sequencing of small RNAs. Nucleic Acids Research, 50, e2–e2.

67. Ruiz-Reche, A., Srivastava, A., Indi, J.A., de la Rubia, I. and Eyras, E. (2019) ReorientExpress: reference-free orientation of nanopore cDNA reads with deep learning. Genome Biology, 20, 260.

68. Chang, H., Lim, J., Ha, M. and Kim, V.N. (2014) TAIL-seq: Genome-wide Determination of Poly(A) Tail Length and 3Ͱ End Modifications. Molecular Cell, 53, 1044–1052.

69. Nicholson, A.L. and Pasquinelli, A.E. (2019) Tales of Detailed Poly(A) Tails. Trends Cell Biol, 29, 191–200.

70. Beverly, M., Hagen, C. and Slack, O. (2018) Poly A tail length analysis of in vitro transcribed mRNA by LC-MS. Anal Bioanal Chem, 410, 1667–1677.

71. Heinicke, F., Zhong, X., Zucknick, M., Breidenbach, J., Sundaram, A.Y.M., T. Flåm, S., Leithaug, M., Dalland, M., Farmer, A., Henderson, J.M., et al. (2020) Systematic assessment of commercially available low-input miRNA library preparation kits. RNA Biology, 17, 75–86.

72. Workman, R.E., Tang, A.D., Tang, P.S., Jain, M., Tyson, J.R., Razaghi, R., Zuzarte, P.C., Gilpatrick, T., Payne, A., Quick, J., et al. (2019) Nanopore native RNA sequencing of a human poly(A) transcriptome. Nat Methods, 16, 1297–1305.

73. Krause, M., Niazi, A.M., Labun, K., Cleuren, Y.N.T., Müller, F.S. and Valen, E. (2019) tailfindr: alignment-free poly(A) length measurement for Oxford Nanopore RNA and DNA sequencing. RNA, 25, 1229–1241.

74. Krogh, A. (1998) Chapter 4 - An introduction to hidden Markov models for biological sequences. In Salzberg, S.L., Searls, D.B., Kasif, S. (eds), New Comprehensive Biochemistry, Computational Methods in Molecular Biology. Elsevier, Vol. 32, pp. 45–63.

75. Edmonds, M. and Winters, M.A. (1976) Polyadenylate Polymerases. In Cohn, W.E. (ed), Progress in Nucleic Acid Research and Molecular Biology. Academic Press, Vol. 17, pp. 149–179.

76. Cao, G.J. and Sarkar, N. (1992) Identification of the gene for an Escherichia coli poly(A) polymerase. Proceedings of the National Academy of Sciences, 89, 10380–10384.

77. Sippel, A.E. (1973) Purification and Characterization of Adenosine Triphosphate: Ribonucleic Acid Adenyltransferase from Escherichia coli. European Journal of Biochemistry, 37, 31–40.

78. Yehudai-Resheff, S. and Schuster, G. (2000) Characterization of the E.coli poly(A) polymerase: nucleotide specificity, RNA-binding affinities and RNA structure dependence. Nucleic Acids Research, 28, 1139–1144.

79. Martin, G. and Keller, W. (1998) Tailing and 3’-end labeling of RNA with yeast poly(A) polymerase and various nucleotides. RNA, 4, 226–230.

80. Sarkar, N. (1997) POLYADENYLATION OF mRNA IN PROKARYOTES. Annu. Rev. Biochem., 66, 173–197.

81. Van Goethem, A., Yigit, N., Everaert, C., Moreno-Smith, M., Mus, L.M., Barbieri, E., Speleman, F., Mestdagh, P., Shohet, J., Van Maerken, T., et al. (2016) Depletion of tRNA-halves enables effective small RNA sequencing of low-input murine serum samples. Sci Rep, 6, 37876.

82. Haile, S., Corbett, R.D., Bilobram, S., Mungall, K., Grande, B.M., Kirk, H., Pandoh, P., MacLeod, T., McDonald, H., Bala, M., et al. (2019) Evaluation of protocols for rRNA depletion-based RNA sequencing of nanogram inputs of mammalian total RNA. PLOS ONE, 14, e0224578.

83. Phelps, W.A., Carlson, A.E. and Lee, M.T. (2021) Optimized design of antisense oligomers for targeted rRNA depletion. Nucleic Acids Research, 49, e5.

84. Loi, D.S.C., Yu, L. and Wu, A.R. (2021) Effective ribosomal RNA depletion for single-cell total RNA-seq by scDASH. PeerJ, 9, e10717.

85. Gu, W., Crawford, E.D., O’Donovan, B.D., Wilson, M.R., Chow, E.D., Retallack, H. and DeRisi, J.L. (2016) Depletion of Abundant Sequences by Hybridization (DASH): using Cas9 to remove unwanted high-abundance species in sequencing libraries and molecular counting applications. Genome Biology, 17, 41.

86. Kraus, A.J., Brink, B.G. and Siegel, T.N. (2019) Efficient and specific oligo-based depletion of rRNA. Sci Rep, 9, 12281.

87. Xiao, L., Wang, J., Ju, S., Cui, M. and Jing, R. (2022) Disorders and roles of tsRNA, snoRNA, snRNA and piRNA in cancer. Journal of Medical Genetics, 59, 623–631.

88. Balatti, V., Nigita, G., Veneziano, D., Drusco, A., Stein, G.S., Messier, T.L., Farina, N.H., Lian, J.B., Tomasello, L., Liu, C., et al. (2017) tsRNA signatures in cancer. Proceedings of the National Academy of Sciences, 114, 8071–8076.

89. Fu, B.-F. and Xu, C.-Y. (2022) Transfer RNA-Derived Small RNAs: Novel Regulators and Biomarkers of Cancers. Frontiers in Oncology, 12.

90. Liu, B., Cao, J., Wang, X., Guo, C., Liu, Y. and Wang, T. (2021) Deciphering the tRNA-derived small RNAs: origin, development, and future. Cell Death Dis, 13, 1–13.

91. DiStefano, J.K. (2018) The Emerging Role of Long Noncoding RNAs in Human Disease. Methods Mol Biol, 1706, 91–110.

92. Hücker, S.M., Fehlmann, T., Werno, C., Weidele, K., Lüke, F., Schlenska-Lange, A., Klein, C.A., Keller, A. and Kirsch, S. (2021) Single-cell microRNA sequencing method comparison and application to cell lines and circulating lung tumor cells. Nat Commun, 12, 4316.

93. Poptsova, M.S., Il’icheva, I.A., Nechipurenko, D.Y., Panchenko, L.A., Khodikov, M.V., Oparina, N.Y., Polozov, R.V., Nechipurenko, Y.D. and Grokhovsky, S.L. (2014) Non-random DNA fragmentation in next-generation sequencing. Sci Rep, 4, 4532.

94. Love, M.I., Hogenesch, J.B. and Irizarry, R.A. (2016) Modeling of RNA-seq fragment sequence bias reduces systematic errors in transcript abundance estimation. Nat Biotechnol, 34, 1287–1291.

95. Oshlack, A. and Wakefield, M.J. (2009) Transcript length bias in RNA-seq data confounds systems biology. Biology Direct, 4, 14.

96. Mandelboum, S., Manber, Z., Elroy-Stein, O. and Elkon, R. (2019) Recurrent functional misinterpretation of RNA-seq data caused by sample-specific gene length bias. PLOS Biology, 17, e3000481.

97. Gorin, G. and Pachter, L. (2021) Length Biases in Single-Cell RNA Sequencing of pre-mRNA. 10.1101/2021.07.30.454514.

98. Gohl, D.M., Magli, A., Garbe, J., Becker, A., Johnson, D.M., Anderson, S., Auch, B., Billstein, B., Froehling, E., McDevitt, S.L., et al. (2019) Measuring sequencer size bias using REcount: a novel method for highly accurate Illumina sequencing-based quantification. Genome Biology, 20, 85.

99. Dmitriev, P., Barat, A., Polesskaya, A., O’Connell, M.J., Robert, T., Dessen, P., Walsh, T.A., Lazar, V., Turki, A., Carnac, G., et al. (2013) Simultaneous miRNA and mRNA transcriptome profiling of human myoblasts reveals a novel set of myogenic differentiation-associated miRNAs and their target genes. BMC Genomics, 14, 265.

100. Pagès, H., Aboyoun, P., Gentleman, R. and DebRoy, S. (2020) Biostrings: Efficient manipulation of biological strings.

