## Supplemental Methods and Results for "To make a short story long: simultaneous short and long RNA profiling on Nanopore devices"

### Supplementary Material and Methods

#### Protocol for the construction of equimolar microRNA HiM and LiM mixes

1. Pipette 1 microliter from each of the indicated diluted 1:10 stock microRNAs into a 200 microliter Eppendorf tube labelled as **HiM**
2. Pipette 95 microliters of RNAse free water into the Eppendorf tube labelled as **HiMs(tock).** The expected concentration (total) of the microRNAs is 50 pmole/100 μL = 0.5 pmole/μL = 500 fmole/μL.
3. Take 1 μL of the **HiMs** and mix it with 19 μL of RNAse free water to construct the **HiM** solution. The expected total concentration is 500/20 = 25 fmole/μL
4. Pipette 1 microliter from each of the indicated diluted stock microRNAs into a 200 microliter Eppendorf tube labelled as **LiM**
5. Pipette 195 microliter of RNAse free water into the Eppendorf tube labelled as **LiMs(tock).** The expected concentration (total) of microRNAs in 50 pmole/200μL = 250 fmole/μL.
6. Take 1 μL of the **LiMs** and mix it with 19 μL of RNAse free water to construct the **LiM** solution. The expected total concentration is 250/20 = 12.5 fmole/μL

#### Construction of synthetic mixes of short and long RNAs

One microliter from the stock solution of each microRNA was mixed with 9 μL of RNAse free water to obtain 1:10 diluted stock solutions of each of the ten stocks immediately prior to the construction of synthetic RNA libraries. While the theoretical concentration of each of the ten working solutions is 10 μΜ, pipetting errors and variable rates of degradation of the RNA in storage will make the actual concentrations to deviate from the expected one. Therefore, the concentrations of RNAs in the working solutions were measured in the Qubit flurometer prior to construction of libraries that used these RNAs. miRNAs were quantified according to the formula:

$$fmoles ssRNA/\mu L= \frac{mass of ssRNA \left( ng/\mu L \right)\times1,000,000}{length of ssRNA \left( nt \right)\times321.47+18.02}$$

The ten microRNAs were subsequently pooled into two groups: a high concentration “**HiMs**” and a low concentration “**LiMs**” equimolar microRNA stock solutions in which the theoretical concentration of each of the HiMs microRNAs was double the concentration of the LiMs group (**Supplementary Methods**). These stock solutions were further diluted 1:20 to derive working **HiM** and **LiM** solutions. . A *working ERCC* (1:20) *solution* was constructed by combining 1 μL of the ERCC mix with 19 μL of RNAse free water for a final concentration of ~ 5 fmol/μL. This working solution was used as input for the construction of samples that were subjected to sequencing. The following table shows the amount of miRNA and ERCC working solutions that need to be combined to derive the synthetic mixes used for subsequent analyses. For the experiments simulating a short RNA library the stock microRNA solutions (HIMs and LiMs) were used as input, while for the experiments simulating simultaneous detection of short and long RNAs, the working microRNA solutions (HiM and LiM) were used to make libraries.

| **Synthetic mix** | **Volume of LMi Solution** | **Volume of HMi Solution** | **Volume of ERCC working solution** | **Total Volume** |
| --- | --- | --- | --- | --- |
| **LMi+HMi+ERCC** | 1 μL | 1 μL | 2 μL | 4 μL |
| **ERCC** | 0 μL | 0 μL | 2 μL | 2 μL |

#### Poly-Adenylation Lengthening of Short RNAs for Simultaneous Short and Long Nanopore Sequencing (PALS-NS) Protocol

The following is an example protocol assuming one is running four libraries at a time. Unless otherwise stated, volumes listed are volumes per library.

A. **Polyadenylation**
1. At room temperature, add the tailing reagents in the order shown in a PCR tube:

Modified poly-A tailing protocol reaction mix

| **Amount** | **Component** |
| --- | --- |
| X μL | Library^1^ |
| 6 – X μL | Nuclease-free Water |
| 1 μL | 10 x *E*-PAP Buffer^2^ |
| 1 μL | 10 mM ATP |
| 1 μL | PAP (4 units)^3^ |
| 9µL | Final Reaction Volume |

^1^The library may contain an arbitrary number of fmoles of RNA input.

^2^We use a polyA Polymerase(PAP) in a MgCl2 reaction buffer e.g. Lucigen’s Cat. No. PAP5104H.

^3^The amount PAP shown above should be sufficient for inputs between 5-1200 fmoles.

If one decides to incorporate a ERCC spike-in to a biological sample of interest, we suggest the following poly-A protocol

| **Amount** | **Component** |
| --- | --- |
| X μL | Library |
| 5 – X μL | Nuclease-free Water |
| 1 μL | 1:100 ERCC mix |
| 1 μL | 10 x *E*-PAP Buffer |
| 1 μL | 10 mM ATP |
| 1 μL | PAP (4 units) |
| 9µL | Final Reaction Volume |

2. Incubate the four reactions at 37°C for 15 minutes.
3. Stop the reaction by freezing immediately to -20oC for 10 minutes

**OPTIONAL STOPPING POINT (samples should be kept at -20oC)**

B. **Reverse Transcription and Strand Switching**
4. In the **PCR tube used to carry out the poly-adenylation** reaction add:
   a. 1µL of VN Primers at 2µM (found in the nanopore kit).
   b. 1µL of 10mM dNTP
5. Mix gently by flicking the tube, and spin down.
6. Incubate at 65°C for 5 minutes and then snap cool on a pre-chilled freezer block.
7. In a separate tube, mix together the following:
   c. 4µL 5x RT Buffer
   d. 1µL RNaseOUT
   e. 1µL Nuclease-free water
   f. 2µL Strand-Switching Primer (SSP, at 10µM)
8. Mix gently by flicking the tube and spin down.
9. Add the strand-switching buffer to the snap-cooled, annealed mRNA, mix by flicking the tube and spin down.
10. Incubate at 42°C for 2 minutes.
11. Add 1µL of Maxima H Minus Reverse Transcriptase. The total volume is now 20µL.
12. Mix gently by flicking the tube and spin down.
13. Incubate the reactions using the following protocol:
    g. Reverse transcription and strand-switching 90 minutes at 42°C (1 cycle).
    h. Heat inactivation for 5 minutes at 85°C (1 cycle).
    i. Hold at 4°C

**OPTIONAL STOPPING POINT (samples should be kept at 4°C )**

C. **Selecting Full-Length Transcripts by PCR**
14. Aliquot the 20μL product from each reverse transcript reaction to four PCR tubes
15. For each of the four PCR tubes of step 1, prepare the following reaction at room temperature:
    a. 25µL of 2x LongAmp Taq Master Mix
    b. 1.5µL cDNA Primer (cPRM)
    c. 18.5µL Nuclease-free water
    d. 5µL Reverse-transcribed RNA sample
16. Amplify using the following cycling conditions:
    e. Initial denaturation for 30 seconds at 95°C (1 cycle)
    f. PCR program settings (number of cycles depending on input):
       i. Denaturation for 15 seconds at 95°C
       ii. Annealing for 15 seconds at 62°C
       iii. Extension for **3 - 6 minutes** at 65°C (aim for **50 seconds per kb** so that 3 minutes will amplify the synthetic ERCC sample and 6 minutes will amplify most human RNAs)
    g. Final extension for **3-6** minutes at 65°C (1 cycle)
    h. Hold at 4°C

**OPTIONAL STOPPING POINT (samples could be kept at 4°C )**

17. Add 1µL of NEB Exonuclease 1 directly to each PCR tube. Mix by pipetting.
18. Incubate the reaction at 37°C for 15 minutes, followed by 80°C for 15 minutes.
19. Pool the four PCR reactions (total 204µL) in a clean 1.5mL Eppendorf DNA LoBind tube.
D. **PCR product cleanup**
20. Resuspend the AMPure XP beads by vortexing.
21. Add 368µL of resuspended AMPure XP beads (1.8x) to the reaction and mix by pipetting.
22. Incubate on a Hula mixer (rotator mixer) for 5 minutes at room temperature.
23. Prepare 500µL of fresh 70% ethanol in Nuclease-free water.
24. Spin down the sample and pellet on a magnet. Keep the tube on the magnet, and pipette off the supernatant.
25. Keep the tube on the magnet and wash the beads with 200µL of freshly prepared 70% ethanol without disturbing the pellet. Remove the ethanol using a pipette and discard.
26. Repeat the aforementioned step.
27. Spin down and place the tube back on the magnet. Pipette off any residual ethanol. Allow to dry for ~30 seconds, but do not dry the pellet to the point of cracking.
28. Remove the tube from the magnetic rack and resuspend the pellet in 12µL of Elution Buffer (EB).
29. Incubate at room temperature for 10 minutes.
30. Pellet the beads on the magnet until the eluate is clear and colorless.
31. Remove and retain the 12µL of eluate into a clean 1.5mL Eppendorf DNA LoBind tube.
32. Remove and retain the eluate which contains the cDNA library in a clean 1.5mL Eppendorf DNA LoBind tube.
33. Dispose of the pelleted beads.
E. **Determining the amount of library to load on the Nanopore Flow Cell**
34. Analyze 1µL of the amplified DNA for size, quantity and quality using the Bioanalyzer. **Use either the high sensitivity (high complexity samples) or the broad band (low to moderate complexity) chips based on the expected concentration of the dominant RNA as well as the RNA species of interest in the sample.**
35. Determine how much library to load on the flow cell . The relevant information can be glanced from the Bioanalyzer chip output (an example is shown below before introducing the formula)
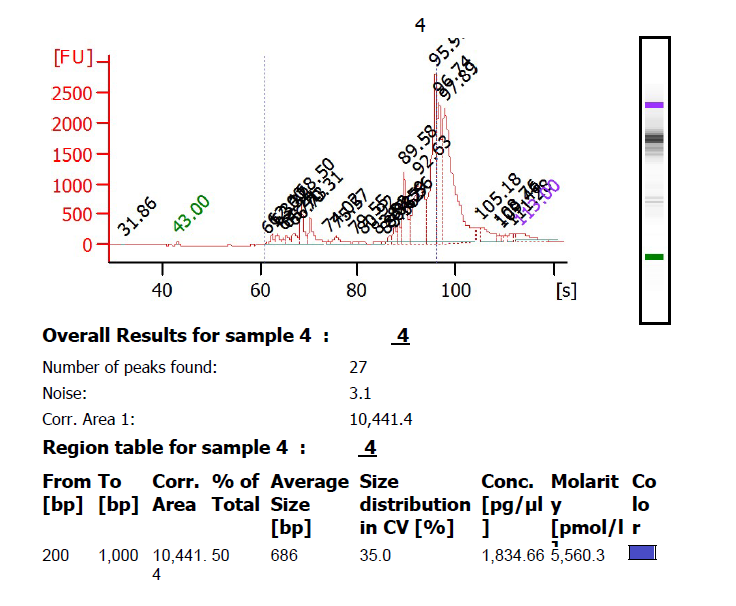

This sample has a concentration of dsDNA that is 5,560.3 pM in the region between 200 to 1000 bases with and average size (in bp) of 686 bp. Therefore, each μL of library contains: 5,560 10^-6^ x pmole = 5,560 x 10^-6^ x 10^3^ pmole i.e 5.56 fmole.

To convert the molarity table for another sample, use the formula:

fmole dsDNA/μL library = Molarity (pmol/l) x 0.001

36. Estimate the amount of library to be carried forward to the sequencing

so that the amount of library that will be used in the next step is:

$$Amount of library (\mu L)=\frac{Target fmole (e.g. 50-200)}{\frac{fmole}{\mu L}dsDNA library}$$

Occasionally the Bioanalyzer will fail to integrate the curve shown above . In such a case, one may input the entire library into the next step, or use volume inputs based on previous runs with the same, or similar samples as rough guides.

F. **Load the Flow Cell and set up the sequencing run as per Oxford Nanopore’s instructions**

### Supplementary Tables

**Supplementary Table 1** Regular expressions used to identify poly-A tails in PALS-NS inserts

| Poly-A pattern | Regular Expression Pattern | Interpretation |
| --- | --- | --- |
| polyAinter0 | A{2,}([CTG]){0,2}([ACTG]{0,8}) | At least two As, optional C/T/G (up to 2), followed by 0-8 A/C/T/G |
| polyAinter1 | A{2,}([CTG]){1,2}A{2,}([CTG]){0,2}([ACTG]{0,8}) | At least two As, followed by one or two C/T/G, followed by two or more As, then 0-2 C/T/G, followed by 0-8 A/C/T/G |
| polyAinter2 | A{2,}([CTG]){1,2}A{2,}([CTG]){1,2}A{2,}([CTG]){0,2}([ACTG]{0,8}) | At least two As, followed by one or two C/T/G, followed by two or more As, then 1-2 C/T/G, followed by two or more As, followed by 0-8 A/C/T/G |

**Supplementary Table 2** MicroRNAs used in sequencing experiments

| microRNA | Accession Number | Sequence | Length (nt) | Group |
| --- | --- | --- | --- | --- |
| hsa-miR-192-5p | MIMAT0000222 | CUGACCUAUGAAUUGACAGCC | 21 | HiM |
| hsa-miR-21-5p | MIMAT0000076 | UAGCUUAUCAGACUGAUGUUGA | 22 | HiM |
| hsa-miR-200b-5p | MIMAT0004571 | CAUCUUACUGGGCAGCAUUGGA | 22 | HiM |
| hsa-miR-200c-5p | MIMAT0004657 | CGUCUUACCCAGCAGUGUUUGG | 22 | HiM |
| hsa-miR-143-5p | MIMAT0004599 | GGUGCAGUGCUGCAUCUCUGGU | 22 | LiM |
| hsa-miR-145-5p | MIMAT0000437 | GUCCAGUUUUCCCAGGAAUCCCU | 23 | HiM |
| hsa-miR-132-5p | MIMAT0004594 | ACCGUGGCUUUCGAUUGUUACU | 22 | LiM |
| hsa-miR-483-3p | MIMAT0002173 | UCACUCCUCUCCUCCCGUCUU | 21 | LiM |
| hsa-miR-744-5p | MIMAT0004945 | UGCGGGGCUAGGGCUAACAGCA | 22 | LiM |
| hsa-miR-324-3p | MIMAT0000762 | CCCACUGCCCCAGGUGCUGCUGG | 23 | LiM |

HiM: High Concentration microRNA, LiM: Low Concentration microRNA

**Supplementary Table 3** RNA characteristics and composition of the ERCC mix

| ERCCID | Length (nt) | MW | Concentration (fmol/μl) | Concentration (ng/μl) | Copy Number |
| --- | --- | --- | --- | --- | --- |
| ERCC-00002 | 1061 | 341162.2 | 15 | 5.117433 | 9.03E+09 |
| ERCC-00003 | 1023 | 327529.6 | 0.9375 | 0.307059 | 564562500 |
| ERCC-00004 | 523 | 167215.6 | 7.5 | 1.254117 | 4516500000 |
| ERCC-00009 | 984 | 316583.8 | 0.9375 | 0.296797313 | 564562500 |
| ERCC-00012 | 994 | 320262.8 | 0.000114441 | 3.67E-05 | 68916.32202 |
| ERCC-00013 | 808 | 261414.6 | 0.000915527 | 0.000239332 | 551330.5641 |
| ERCC-00014 | 1957 | 631409.4 | 0.003662109 | 0.00231229 | 2205322.269 |
| ERCC-00016 | 844 | 271683.8 | 0.000228882 | 6.22E-05 | 137832.644 |
| ERCC-00017 | 1136 | 367042.2 | 0.000114441 | 4.20E-05 | 68916.32202 |
| ERCC-00019 | 644 | 207542.8 | 0.029296875 | 0.006080355 | 17642578.13 |
| ERCC-00022 | 751 | 241178.2 | 0.234375 | 0.056526141 | 141140625 |
| ERCC-00024 | 536 | 173128.2 | 0.000228882 | 3.96E-05 | 137832.644 |
| ERCC-00025 | 1994 | 640940.8 | 0.05859375 | 0.037555125 | 35285156.25 |
| ERCC-00028 | 1130 | 364285 | 0.003662109 | 0.001334052 | 2205322.269 |
| ERCC-00031 | 1138 | 365731.6 | 0.001831055 | 0.000669675 | 1102661.134 |
| ERCC-00033 | 2022 | 651534.4 | 0.001831055 | 0.001192995 | 1102661.134 |
| ERCC-00034 | 1019 | 328138.8 | 0.007324219 | 0.00240336 | 4410644.531 |
| ERCC-00035 | 1130 | 364378 | 0.1171875 | 0.042700547 | 70570312.5 |
| ERCC-00039 | 740 | 238322 | 0.003662109 | 0.000872761 | 2205322.269 |
| ERCC-00040 | 744 | 239737.8 | 0.000915527 | 0.000219487 | 551330.5641 |
| ERCC-00041 | 1122 | 362677.4 | 0.000228882 | 8.30E-05 | 137832.644 |
| ERCC-00042 | 1023 | 325749.6 | 0.46875 | 0.152695125 | 282281250 |
| ERCC-00043 | 1023 | 330121.6 | 0.46875 | 0.1547445 | 282281250 |
| ERCC-00044 | 1156 | 372347.2 | 0.1171875 | 0.043634438 | 70570312.5 |
| ERCC-00046 | 522 | 168087.4 | 3.75 | 0.63032775 | 2258250000 |
| ERCC-00048 | 992 | 320110.4 | 1.43E-05 | 4.58E-06 | 8614.543264 |
| ERCC-00051 | 274 | 88355.8 | 0.05859375 | 0.005177098 | 35285156.25 |
| ERCC-00053 | 1023 | 327970.6 | 0.029296875 | 0.009608514 | 17642578.13 |
| ERCC-00054 | 274 | 88965.8 | 0.014648438 | 0.00130321 | 8821289.063 |
| ERCC-00057 | 1021 | 328287.2 | 1.43E-05 | 4.70E-06 | 8614.543264 |
| ERCC-00058 | 1136 | 366548.2 | 0.001831055 | 0.00067117 | 1102661.134 |
| ERCC-00059 | 525 | 168750 | 0.014648438 | 0.002471924 | 8821289.063 |
| ERCC-00060 | 523 | 168194.6 | 0.234375 | 0.039420609 | 141140625 |
| ERCC-00061 | 1136 | 366454.2 | 5.72E-05 | 2.10E-05 | 34458.16101 |
| ERCC-00062 | 1023 | 328504.6 | 0.05859375 | 0.019248316 | 35285156.25 |
| ERCC-00067 | 644 | 207450.8 | 0.003662109 | 0.000759708 | 2205322.269 |
| ERCC-00069 | 1137 | 366664.4 | 0.001831055 | 0.000671383 | 1102661.134 |
| ERCC-00071 | 642 | 206115.4 | 0.05859375 | 0.012077074 | 35285156.25 |
| ERCC-00073 | 603 | 193957.6 | 0.000915527 | 0.000177573 | 551330.5641 |
| ERCC-00074 | 522 | 167539.4 | 15 | 2.513091 | 9.03E+09 |
| ERCC-00075 | 1023 | 325441.6 | 1.43E-05 | 4.66E-06 | 8614.543264 |
| ERCC-00076 | 642 | 206436.4 | 0.234375 | 0.048383531 | 141140625 |
| ERCC-00077 | 273 | 87693.6 | 0.003662109 | 0.000321144 | 2205322.269 |
| ERCC-00078 | 993 | 320093.6 | 0.029296875 | 0.009377742 | 17642578.13 |
| ERCC-00079 | 644 | 207756.8 | 0.05859375 | 0.01217325 | 35285156.25 |
| ERCC-00081 | 534 | 172322.8 | 0.000228882 | 3.94E-05 | 137832.644 |
| ERCC-00083 | 1022 | 325322.4 | 2.86E-05 | 9.31E-06 | 17229.08051 |
| ERCC-00084 | 994 | 320444.8 | 0.029296875 | 0.009388031 | 17642578.13 |
| ERCC-00085 | 844 | 271322.8 | 0.007324219 | 0.001987228 | 4410644.531 |
| ERCC-00086 | 1020 | 328632 | 0.000114441 | 3.76E-05 | 68916.32202 |
| ERCC-00092 | 1124 | 361715.8 | 0.234375 | 0.084777141 | 141140625 |
| ERCC-00095 | 521 | 166307.2 | 0.1171875 | 0.019489125 | 70570312.5 |
| ERCC-00096 | 1107 | 356565.4 | 15 | 5.348481 | 9.03E+09 |
| ERCC-00097 | 523 | 167188.6 | 0.000457764 | 7.65E-05 | 275665.2821 |
| ERCC-00098 | 1143 | 368969.6 | 5.72E-05 | 2.11E-05 | 34458.16101 |
| ERCC-00099 | 1350 | 434408 | 0.014648438 | 0.006363398 | 8821289.063 |
| ERCC-00104 | 2022 | 647370.4 | 0.000228882 | 0.000148171 | 137832.644 |
| ERCC-00108 | 1022 | 328424.4 | 0.9375 | 0.307897875 | 564562500 |
| ERCC-00109 | 536 | 172925.2 | 0.000915527 | 0.000158318 | 551330.5641 |
| ERCC-00111 | 994 | 319358.8 | 0.46875 | 0.149699438 | 282281250 |
| ERCC-00112 | 1136 | 364932.2 | 0.1171875 | 0.042765492 | 70570312.5 |
| ERCC-00113 | 840 | 270697 | 3.75 | 1.01511375 | 2258250000 |
| ERCC-00116 | 1991 | 639986.2 | 0.46875 | 0.299993531 | 282281250 |
| ERCC-00117 | 1136 | 365757.2 | 5.72E-05 | 2.09E-05 | 34458.16101 |
| ERCC-00120 | 536 | 172605.2 | 0.000915527 | 0.000158025 | 551330.5641 |
| ERCC-00123 | 1022 | 324911.4 | 0.000228882 | 7.44E-05 | 137832.644 |
| ERCC-00126 | 1118 | 359444.6 | 0.014648438 | 0.005265302 | 8821289.063 |
| ERCC-00130 | 1059 | 342267.8 | 30 | 10.268034 | 1.81E+10 |
| ERCC-00131 | 771 | 248276.2 | 0.1171875 | 0.029094867 | 70570312.5 |
| ERCC-00134 | 274 | 88593.8 | 0.001831055 | 0.00016222 | 1102661.134 |
| ERCC-00136 | 1033 | 333362.6 | 1.875 | 0.625054875 | 1129125000 |
| ERCC-00137 | 537 | 173218.4 | 0.000915527 | 0.000158586 | 551330.5641 |
| ERCC-00138 | 1024 | 328254.6 | 0.000114441 | 3.76E-05 | 68916.32202 |
| ERCC-00142 | 493 | 159089.6 | 0.000228882 | 3.64E-05 | 137832.644 |
| ERCC-00143 | 784 | 251704.8 | 0.003662109 | 0.000921771 | 2205322.269 |
| ERCC-00144 | 538 | 173403.6 | 0.029296875 | 0.005080184 | 17642578.13 |
| ERCC-00145 | 1042 | 336179.4 | 0.9375 | 0.315168188 | 564562500 |
| ERCC-00147 | 1023 | 331124.6 | 0.000915527 | 0.000303154 | 551330.5641 |
| ERCC-00148 | 494 | 159910.8 | 0.014648438 | 0.002342443 | 8821289.063 |
| ERCC-00150 | 743 | 239127.6 | 0.003662109 | 0.000875711 | 2205322.269 |
| ERCC-00154 | 537 | 173317.4 | 0.007324219 | 0.001269415 | 4410644.531 |
| ERCC-00156 | 494 | 159198.8 | 0.000457764 | 7.29E-05 | 275665.2821 |
| ERCC-00157 | 1019 | 328634.8 | 0.007324219 | 0.002406993 | 4410644.531 |
| ERCC-00158 | 1027 | 330733.4 | 0.000457764 | 0.000151398 | 275665.2821 |
| ERCC-00160 | 743 | 239436.6 | 0.007324219 | 0.001753686 | 4410644.531 |
| ERCC-00162 | 523 | 166408.6 | 0.05859375 | 0.009750504 | 35285156.25 |
| ERCC-00163 | 543 | 174948.6 | 0.014648438 | 0.002562724 | 8821289.063 |
| ERCC-00164 | 1022 | 324758.4 | 0.000457764 | 0.000148663 | 275665.2821 |
| ERCC-00165 | 872 | 279788.4 | 0.05859375 | 0.016393852 | 35285156.25 |
| ERCC-00168 | 1024 | 326398.8 | 0.000457764 | 0.000149414 | 275665.2821 |
| ERCC-00170 | 1023 | 330462.6 | 0.014648438 | 0.004840761 | 8821289.063 |
| ERCC-00171 | 505 | 163022 | 3.75 | 0.6113325 | 2258250000 |
| Total per μL |  |  | 103.52 | 29.97 | 62336750436 |

**Supplementary Table 4** Alignment of microRNAs to the ERCC RNA pool

| miRNA sequence | ERCC sequence | Alignment Score | Number of matches | Number of mismatches | Alignment Start | ERCC sequence Matched | miRNA sequence matched |
| --- | --- | --- | --- | --- | --- | --- | --- |
| hsa-miR-192-5p | ERCC-00024 | 17.836 | 9 | 0 | 308 | GAATTGACA | GAATTGACA |
| hsa-miR-21-5p | ERCC-00069 | 17.836 | 9 | 0 | 15 | TATCAGACT | TATCAGACT |
| hsa-miR-200b-5p | ERCC-00147 | 15.900 | 11 | 1 | 677 | ATCTTACTCGGC | ATCTTACTGGGC |
| hsa-miR-200c-5p | ERCC-00034 | 19.818 | 10 | 0 | 5 | TTACCCAGCA | TTACCCAGCA |
| hsa-miR-143-5p | ERCC-00085 | 19.818 | 10 | 0 | 738 | AGTGCTGCAT | AGTGCTGCAT |
| hsa-miR-132-5p | ERCC-00048 | 19.818 | 10 | 0 | 534 | GTGGCTTTCG | GTGGCTTTCG |
| hsa-miR-145-5p | ERCC-00168 | 19.818 | 10 | 0 | 461 | TCCAGTTTTC | TCCAGTTTTC |
| hsa-miR-483-3p | ERCC-00123 | 19.818 | 10 | 0 | 205 | TCTCCTCCCG | TCTCCTCCCG |
| hsa-miR-744-5p | ERCC-00019 | 17.836 | 9 | 0 | 429 | GGGGCTAGG | GGGGCTAGG |
| hsa-miR-324-3p | ERCC-00003 | 19.818 | 10 | 0 | 261 | CACTGCCCCA | CACTGCCCCA |

Pairwise, local (Smith Waterman) alignment with a gap opening and a gap extension penalty of ten and four respectively, using the function “pairwise” alignment in the Bioconductor package Biostrings(100).

**Supplementary Table 5** Synthetic libraries (RNA input), design (replicates) and PCR conditions (cycles, extension time)

| Experiment | RNA targeted | Library Type | PAP | ERCC Source | Library  Replicate | Sequencing  Replicate | Dilution | MiRNA Input (fmoles) | ERCC Input (fmoles) | Total  Input (fmoles) | PCR Cycles | Extension  Time (min) |
| --- | --- | --- | --- | --- | --- | --- | --- | --- | --- | --- | --- | --- |
| 1^st^ 2x2 | Short | LiM+HiM+ERCC | + | A | 1 | 1 | 1 | 1270.289 | 10.4 | 1280.689 | 15 | 6 |
| 2^nd^ 2x2 | Short | LiM+HiM+ERCC | + | A | 2 | 1 | 1 | 1270.289 | 10.4 | 1280.689 | 15 | 6 |
| 1^st^ 2x2 | Long | LiM+HiM+ERCC | - | A | 1 | 1 | 1 | 1270.289 | 10.4 | 1280.689 | 15 | 6 |
| 2^nd^ 2x2 | Long | LiM+HiM+ERCC | - | A | 2 | 1 | 1 | 1270.289 | 10.4 | 1280.689 | 15 | 6 |
| 2^nd^ 2x2 | Long | LiM+HiM+ERCC | - | A | 2 | 2 | 1 | 1270.289 | 10.4 | 1280.689 | 15 | 6 |
| 1^st^ 2x2 | Long | ERCC | + | A | 1 | 1 | 1 | 0 | 10.4 | 10.4 | 15 | 6 |
| 2^nd^ 2x2 | Long | ERCC | + | A | 2 | 2 | 1 | 0 | 10.4 | 10.4 | 15 | 6 |
| 1^st^ 2x2 | Long | ERCC | - | A | 1 | 1 | 1 | 0 | 10.4 | 10.4 | 15 | 6 |
| 2^nd^ 2x2 | Long | ERCC | - | A | 2 | 1 | 1 | 0 | 10.4 | 10.4 | 15 | 3 |
| 2^nd^ 2x2 | Long | ERCC | - | A | 2 | 2 | 1 | 0 | 10.4 | 10.4 | 15 | 3 |
| DS | Long | ERCC | + | B | 1 | 1 | 1 | 0 | 10.4 | 10.4 | 15 | 3 |
| DS | Long | ERCC | + | B | 2 | 1 | 1 | 0 | 10.4 | 10.4 | 15 | 3 |
| DS | Short+Long | LiM+HiM+ERCC | + | B | 1 | 1 | 1 | 47.2 | 10.4 | 57.6 | 15 | 3 |
| DS | Short+Long | LiM+HiM+ERCC | + | B | 1 | 2 | 1 | 47.2 | 10.4 | 57.6 | 15 | 3 |
| DS | Short+Long | LiM+HiM+ERCC | + | B | 1 | 1 | 10 | 4.7 | 1 | 5.7 | 15 | 3 |
| DS | Short+Long | LiM+HiM+ERCC | + | B | 2 | 1 | 10 | 4.7 | 1 | 5.7 | 15 | 3 |
| DS | Short+Long | LiM+HiM+ERCC | + | B | 1 | 1 | 100 | 0.47 | 0.1 | 0.57 | 19 | 3 |
| DS | Short+Long | LiM+HiM+ERCC | + | B | 2 | 1 | 100 | 0.47 | 0.1 | 0.57 | 19 | 3 |
| DS | Short+Long | LiM+HiM+ERCC | + | B | 1 | 1 | 1000 | 0.047 | 0.01 | 0.057 | 23 | 3 |
| DS | Short+Long | LiM+HiM+ERCC | + | B | 2 | 1 | 1000 | 0.047 | 0.01 | 0.057 | 23 | 3 |

DS: Dilution Series, ERCC: External RNA Controls Consortium reference RNA, FLO-MIN106: MinIon Flow Cells, FLO-FLG001 : Flongle Flow Cells, LiM: miRNAs input at lower ratio, HiM: miRNAs input at higher ratio. Estimated input in the DS was 3.3ng (1:1), 333 pg (1:10), 33.3 pg (1:100), 3.3 pg (1:1000).

**Supplementary Table 6** Unique Ensembl gene biotypes detected by each of the three sequencing platforms: Illumina, ONT without PAP and PALS-NS as a proportion of the reads in each of these libraries.

| Gene Biotype | Illumina | | ONT PAP (-) | | PALS-NS | |
| --- | --- | --- | --- | --- | --- | --- |
|  | Control | High Fructose | Control | High Fructose | Control | High Fructose |
| Protein Coding | 68.042 | 68.392 | 74.466 | 71.879 | 72.339 | 69.310 |
| LncRNA | 13.960 | 14.207 | 13.885 | 15.594 | 13.938 | 15.008 |
| Processed Pseudogene | 7.870 | 7.512 | 8.133 | 8.457 | 5.692 | 7.047 |
| TEC | 5.336 | 5.202 | 0.000 | 0.000 | 0.000 | 0.000 |
| Unprocessed Pseudogene | 0.791 | 0.837 | 0.594 | 0.684 | 0.555 | 0.675 |
| MiRNA | 0.647 | 0.602 | 0.552 | 0.693 | 0.992 | 1.061 |
| IG V Gene | 0.602 | 0.664 | 0.271 | 0.307 | 0.178 | 0.266 |
| SnoRNA | 0.597 | 0.524 | 0.182 | 0.354 | 1.736 | 1.639 |
| Transcribed Processed Pseudogene | 0.478 | 0.499 | 0.662 | 0.613 | 0.609 | 0.643 |
| Transcribed Unprocessed Pseudogene | 0.396 | 0.396 | 0.401 | 0.467 | 0.399 | 0.418 |
| SnRNA | 0.391 | 0.342 | 0.193 | 0.335 | 1.698 | 1.914 |
| TR V Gene | 0.227 | 0.190 | 0.016 | 0.005 | 0.011 | 0.014 |
| TR J Gene | 0.128 | 0.116 | 0.016 | 0.000 | 0.000 | 0.000 |
| Pseudogene | 0.074 | 0.074 | 0.089 | 0.085 | 0.075 | 0.106 |
| Mt tRNA | 0.070 | 0.066 | 0.115 | 0.104 | 0.119 | 0.101 |
| Transcribed Unitary Pseudogene | 0.054 | 0.050 | 0.063 | 0.061 | 0.054 | 0.073 |
| IG C Gene | 0.054 | 0.050 | 0.036 | 0.038 | 0.043 | 0.041 |
| IG V Pseudogene | 0.054 | 0.050 | 0.005 | 0.005 | 0.011 | 0.023 |
| ScaRNA | 0.049 | 0.058 | 0.010 | 0.024 | 0.124 | 0.119 |
| IG J Gene | 0.049 | 0.050 | 0.005 | 0.000 | 0.016 | 0.014 |
| TR C Gene | 0.033 | 0.029 | 0.036 | 0.033 | 0.027 | 0.023 |
| Unitary Pseudogene | 0.029 | 0.029 | 0.010 | 0.014 | 0.016 | 0.041 |
| Misc RNA | 0.012 | 0.012 | 0.172 | 0.156 | 0.340 | 0.399 |
| TR J Pseudogene | 0.012 | 0.008 | 0.000 | 0.000 | 0.000 | 0.000 |
| Mt rRNA | 0.008 | 0.008 | 0.010 | 0.009 | 0.011 | 0.009 |
| rRNA | 0.008 | 0.004 | 0.052 | 0.061 | 0.965 | 1.005 |
| TR V Pseudogene | 0.008 | 0.000 | 0.010 | 0.005 | 0.005 | 0.005 |
| IG D Gene | 0.004 | 0.012 | 0.000 | 0.000 | 0.000 | 0.000 |
| Ribozyme | 0.004 | 0.004 | 0.010 | 0.014 | 0.038 | 0.037 |
| IG C Pseudogene | 0.004 | 0.004 | 0.000 | 0.000 | 0.000 | 0.000 |
| TR D Gene | 0.004 | 0.004 | 0.000 | 0.000 | 0.000 | 0.000 |
| Translated Unprocessed Pseudogene | 0.004 | 0.004 | 0.000 | 0.000 | 0.000 | 0.000 |
| ScRNA | 0.000 | 0.000 | 0.005 | 0.005 | 0.005 | 0.005 |
| SRNA | 0.000 | 0.000 | 0.000 | 0.000 | 0.005 | 0.005 |
| IG_D_pseudogene | 0.000 | 0.000 | 0.000 | 0.000 | 0.000 | 0.000 |
| IG_LV_gene | 0.000 | 0.000 | 0.000 | 0.000 | 0.000 | 0.000 |
| IG_pseudogene | 0.000 | 0.000 | 0.000 | 0.000 | 0.000 | 0.000 |

**Supplementary Table 7** Counts of Ensembl gene biotypes (library representation) detected by each of the three sequencing platforms: Illumina, ONT without PAP and PALS-NS.

| Gene Biotype | Illumina | | ONT PAP(-) | | PALS-NS | |
| --- | --- | --- | --- | --- | --- | --- |
|  | Control | High Fructose | Control | High Fructose | Control | High Fructose |
| Protein Coding | 97.790 | 98.180 | 84.677 | 82.837 | 29.020 | 30.687 |
| IG C Gene | 0.735 | 0.538 | 1.412 | 0.943 | 0.026 | 0.023 |
| LncRNA | 0.691 | 0.605 | 3.304 | 3.424 | 11.288 | 12.364 |
| Unprocessed Pseudogene | 0.158 | 0.149 | 2.161 | 2.272 | 0.077 | 0.071 |
| Processed Pseudogene | 0.136 | 0.120 | 3.538 | 2.917 | 0.404 | 0.460 |
| IG V Gene | 0.133 | 0.091 | 0.043 | 0.037 | 0.002 | 0.002 |
| Mt rRNA | 0.103 | 0.079 | 1.425 | 4.285 | 0.584 | 0.493 |
| TEC | 0.102 | 0.091 | 0.000 | 0.000 | 0.000 | 0.000 |
| Transcribed Unprocessed Pseudogene | 0.080 | 0.083 | 0.190 | 0.134 | 0.031 | 0.031 |
| Transcribed Processed Pseudogene | 0.022 | 0.022 | 0.632 | 0.705 | 0.047 | 0.048 |
| Transcribed Unitary Pseudogene | 0.012 | 0.012 | 0.028 | 0.030 | 0.006 | 0.007 |
| Mt tRNA | 0.010 | 0.010 | 0.826 | 0.863 | 10.679 | 8.452 |
| Pseudogene | 0.007 | 0.003 | 0.012 | 0.010 | 0.044 | 0.032 |
| TR C Gene | 0.007 | 0.003 | 0.005 | 0.003 | 0.000 | 0.000 |
| MiRNA | 0.004 | 0.004 | 0.729 | 0.231 | 2.171 | 1.940 |
| SnoRNA | 0.003 | 0.003 | 0.135 | 0.097 | 1.064 | 0.829 |
| TR V Gene | 0.002 | 0.001 | 0.001 | 0.000 | 0.000 | 0.000 |
| SnRNA | 0.002 | 0.001 | 0.016 | 0.022 | 0.273 | 0.278 |
| IG J Gene | 0.001 | 0.001 | 0.000 | 0.000 | 0.000 | 0.000 |
| IG V Pseudogene | 0.001 | 0.001 | 0.000 | 0.000 | 0.000 | 0.000 |
| TR J Gene | 0.001 | 0.000 | 0.001 | 0.000 | 0.000 | 0.000 |
| ScaRNA | 0.000 | 0.000 | 0.002 | 0.002 | 0.018 | 0.018 |
| TR D Gene | 0.000 | 0.000 | 0.000 | 0.000 | 0.000 | 0.000 |
| Misc RNA | 0.000 | 0.000 | 0.022 | 0.020 | 2.208 | 1.988 |
| Unitary Pseudogene | 0.000 | 0.000 | 0.001 | 0.000 | 0.000 | 0.000 |
| TR J Pseudogene | 0.000 | 0.000 | 0.000 | 0.000 | 0.000 | 0.000 |
| Ribozyme | 0.000 | 0.000 | 0.002 | 0.001 | 0.086 | 0.059 |
| IG C Pseudogene | 0.000 | 0.000 | 0.000 | 0.000 | 0.000 | 0.000 |
| rRNA | 0.000 | 0.000 | 0.838 | 1.163 | 41.966 | 42.207 |
| TR V Pseudogene | 0.000 | 0.000 | 0.000 | 0.000 | 0.000 | 0.000 |
| IG D Gene | 0.000 | 0.000 | 0.000 | 0.000 | 0.000 | 0.000 |
| Translated Unprocessed Pseudogene | 0.000 | 0.000 | 0.000 | 0.000 | 0.000 | 0.000 |
| ScRNA | 0.000 | 0.000 | 0.002 | 0.002 | 0.005 | 0.007 |
| SRNA | 0.000 | 0.000 | 0.000 | 0.000 | 0.000 | 0.000 |
| IG D Pseudogene | 0.000 | 0.000 | 0.000 | 0.000 | 0.000 | 0.000 |
| IG LV Gene | 0.000 | 0.000 | 0.000 | 0.000 | 0.000 | 0.000 |
| IG Pseudogene | 0.000 | 0.000 | 0.000 | 0.000 | 0.000 | 0.000 |

**Supplementary Table 8** Comparison of Bias Factors for short RNAs in the PALS-NS and the 4N randomized adapter ligation protocol.

| miRNA | PALS-NS | 4N |
| --- | --- | --- |
| hsa-miR-132-5p | -0.092 | -0.037 |
| hsa-miR-145-5p | 0.200 | -0.301 |
| hsa-miR-192-5p | -0.098 | -0.447 |
| hsa-miR-21-5p | 0.091 | -0.655 |
| hsa-miR-324-3p | -1.261 | 0.786 |
| hsa-miR-744-5p | 0.098 | -0.029 |

To generate these estimates the Negative Binomial random effect models were fit to the PAP+ sequencing experiments and the equimolar hand-mixed microRNAs(38). After fitting the random effects of miRNAs that were present in both datasets were extracted and reported in log10 scale.

### Supplementary Figures

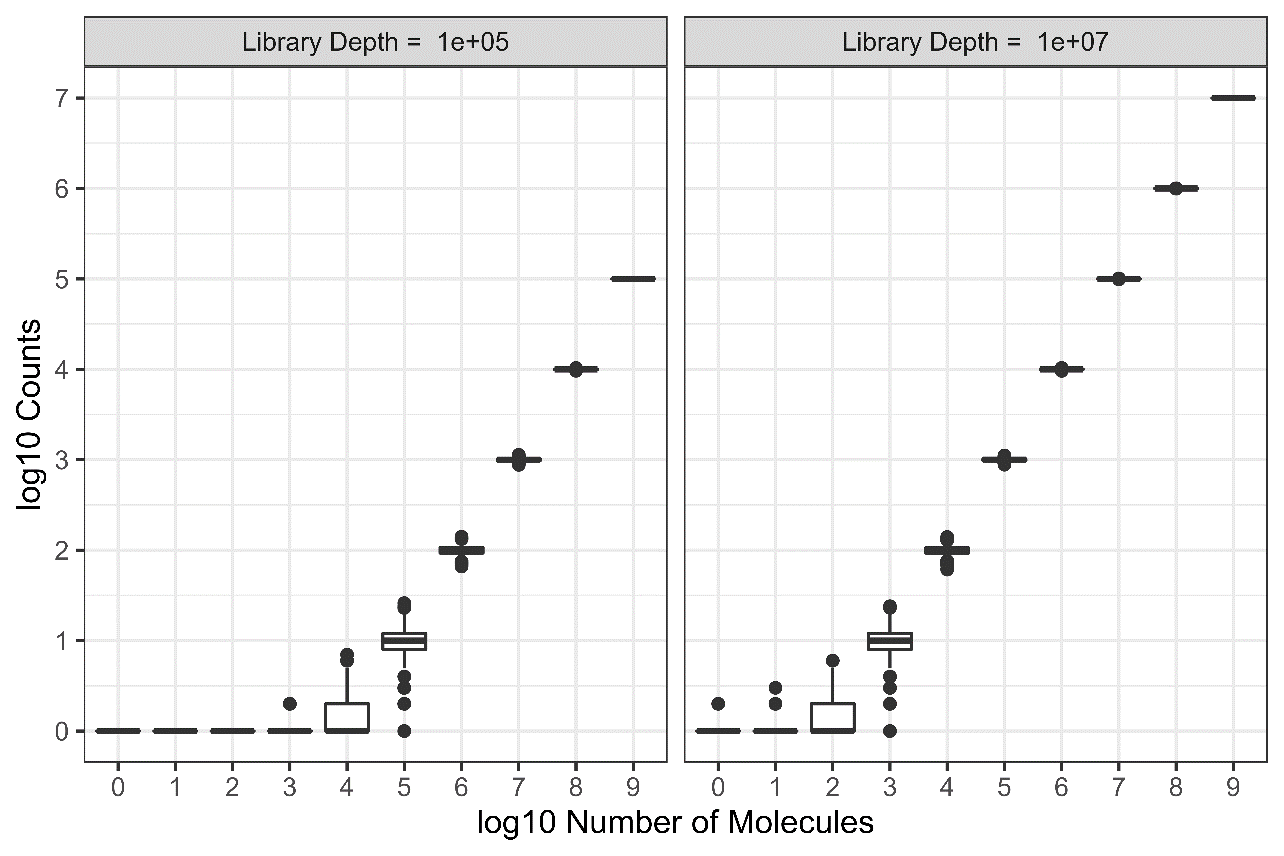

**Supplementary Figure 1** Dynamic range compression during sequencing.

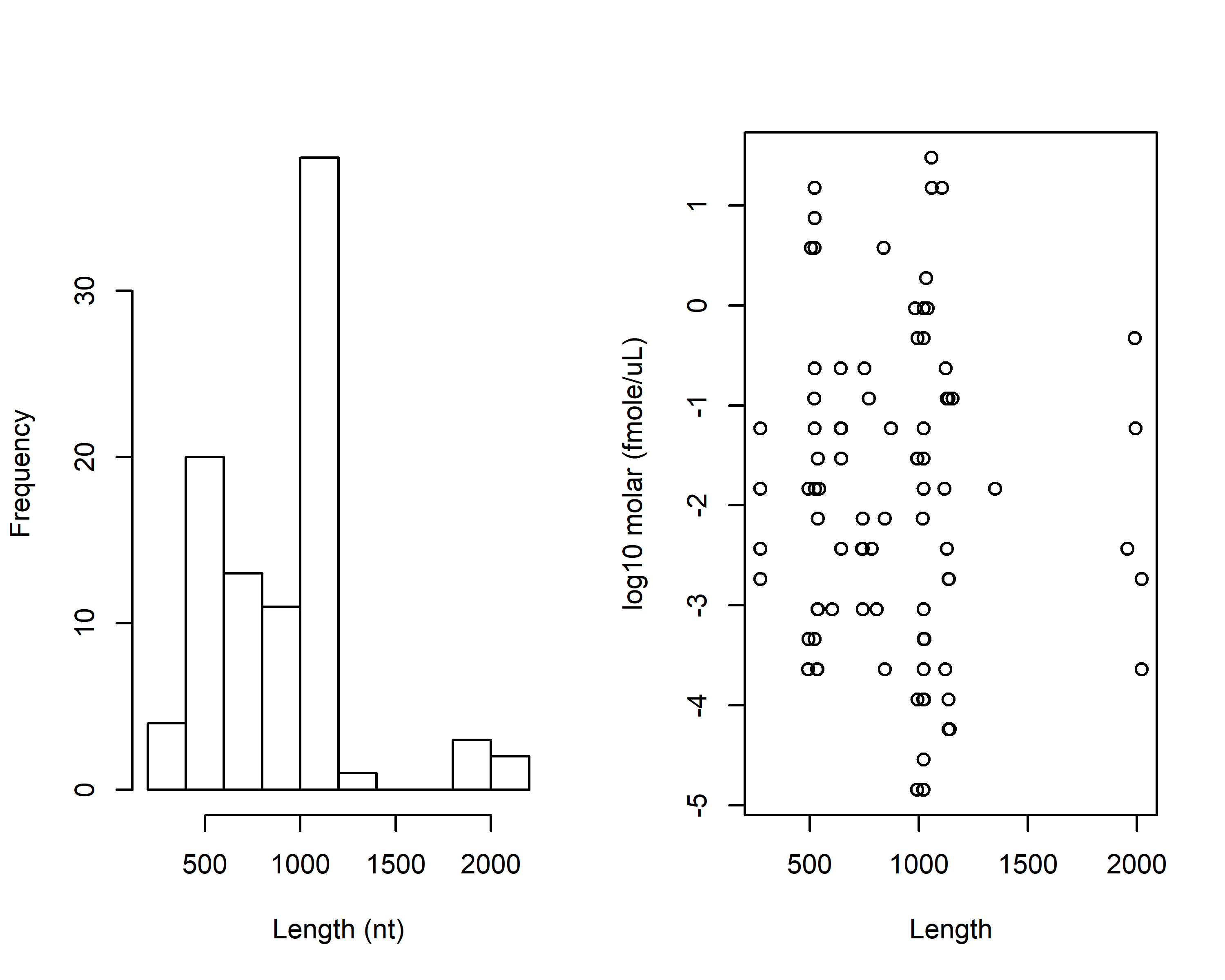

**Supplementary Figure 2** Length distribution and molar amount by sequence length in the ERCC mix.

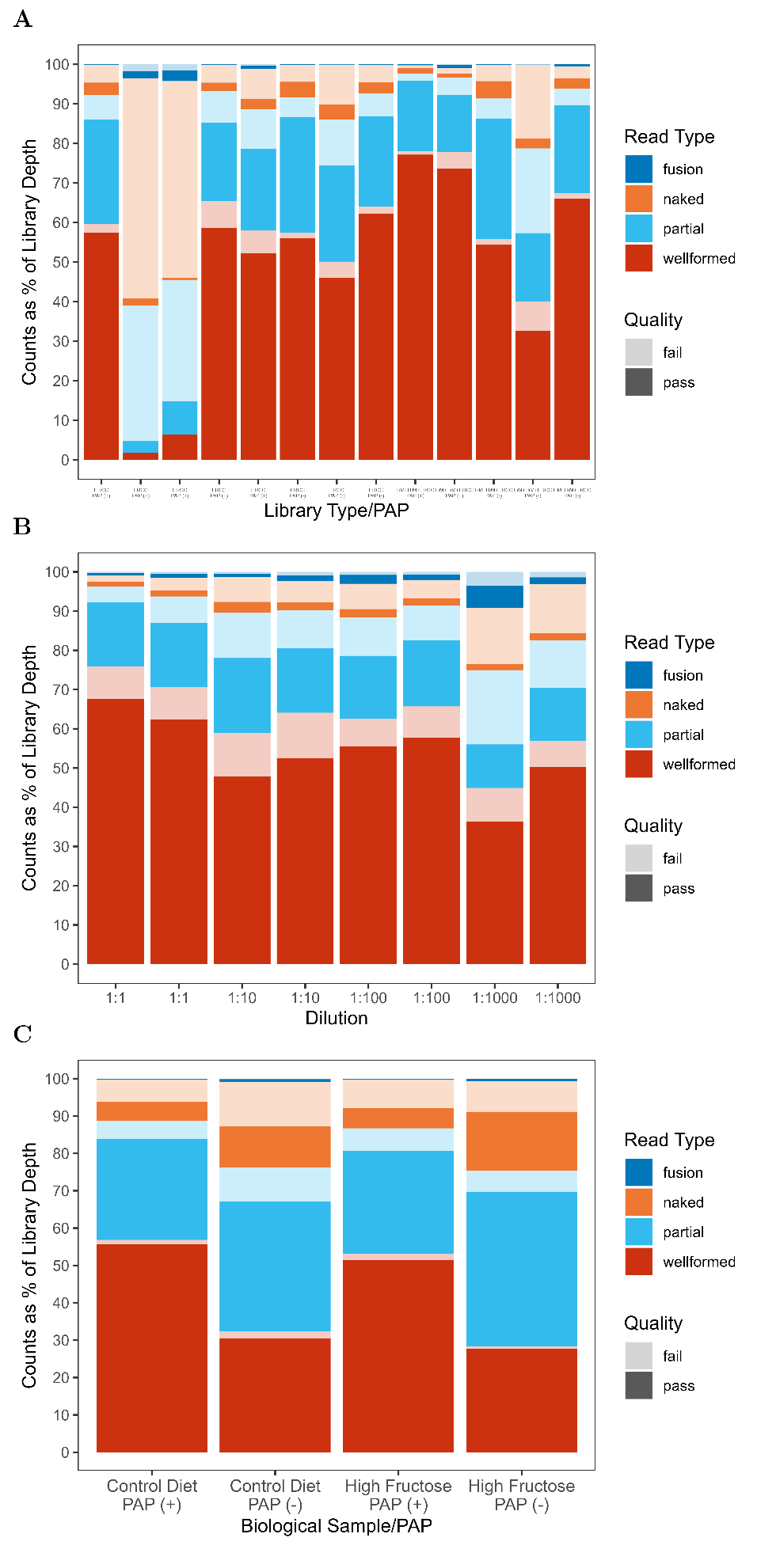

**Supplementary Figure 3** Insert types of sequencing experiments in the 2x2 and the ERCC, non-polyadenylated sample in the dilution series (A), the serial dilutions of the LiM+HiM+ERCC experiments (B) and the biological samples (C). Quality indicated by the transparency scale (opaque for pass, semi-transparent for fail).

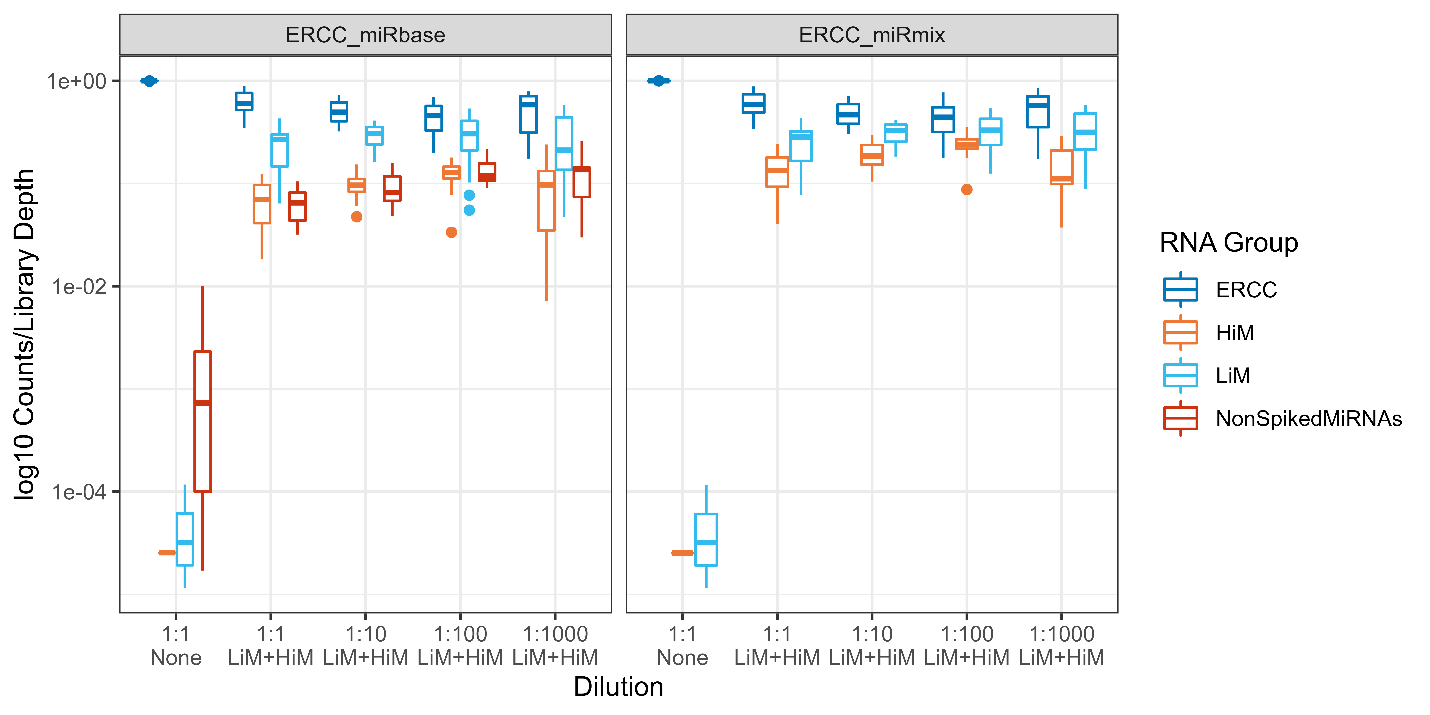

**Supplementary Figure 4** Representation of groups of RNA as a proportion of library depth in the Serial Dilution experiment; for each sub-library in each sequencing run (a total of eight sub-libraries per library) we calculated the representation of RNAs (counts/effective sub-library depth in log10 scale) according to the group they belonged to: ERCC (without any microRNA input, “None”), or ERCC with HiM and LiM (“LiM+HiM”) . Irrespective of the type of microRNA input, all libraries were subjected to poly-adenylation step. We mapped counts against a sequence library that included the 92 ERCC RNAs and the 10 RNAs used in the mix (ERCC_miRmix). A sensitivity analysis was also performed by mapping against a database of the 92 ERCC and the entire miRbase. In the latter case, counts to RNAs not present in the mix (“NonSpikedMiRNAs”) were tabulated as a separate category.

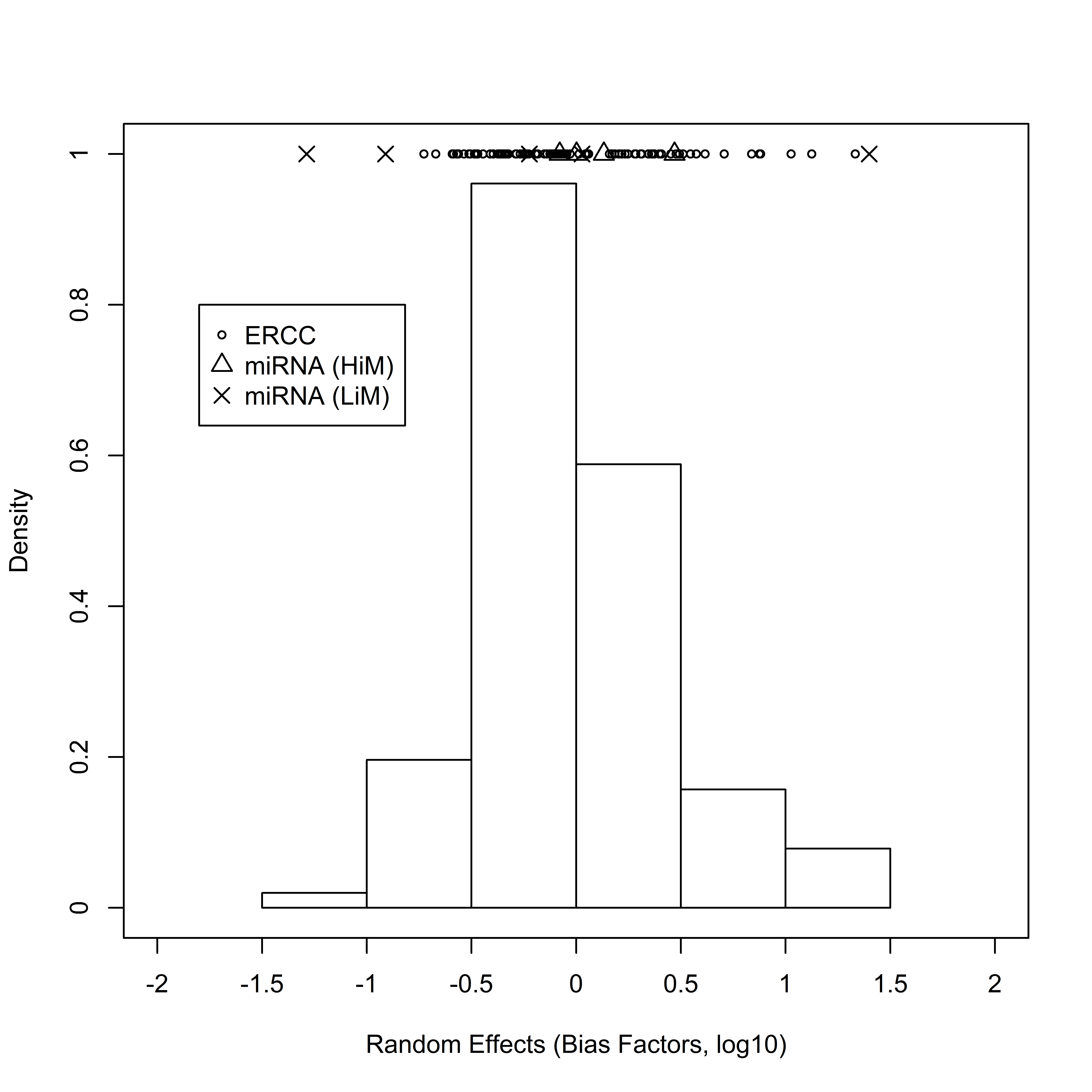

**Supplementary Figure 5** Estimated bias factors of ERCC, HiM and LiM RNAs from the 2x2 and DS experiments. The individual random effects are shown on the top of the graph, along with a histogram of their values.

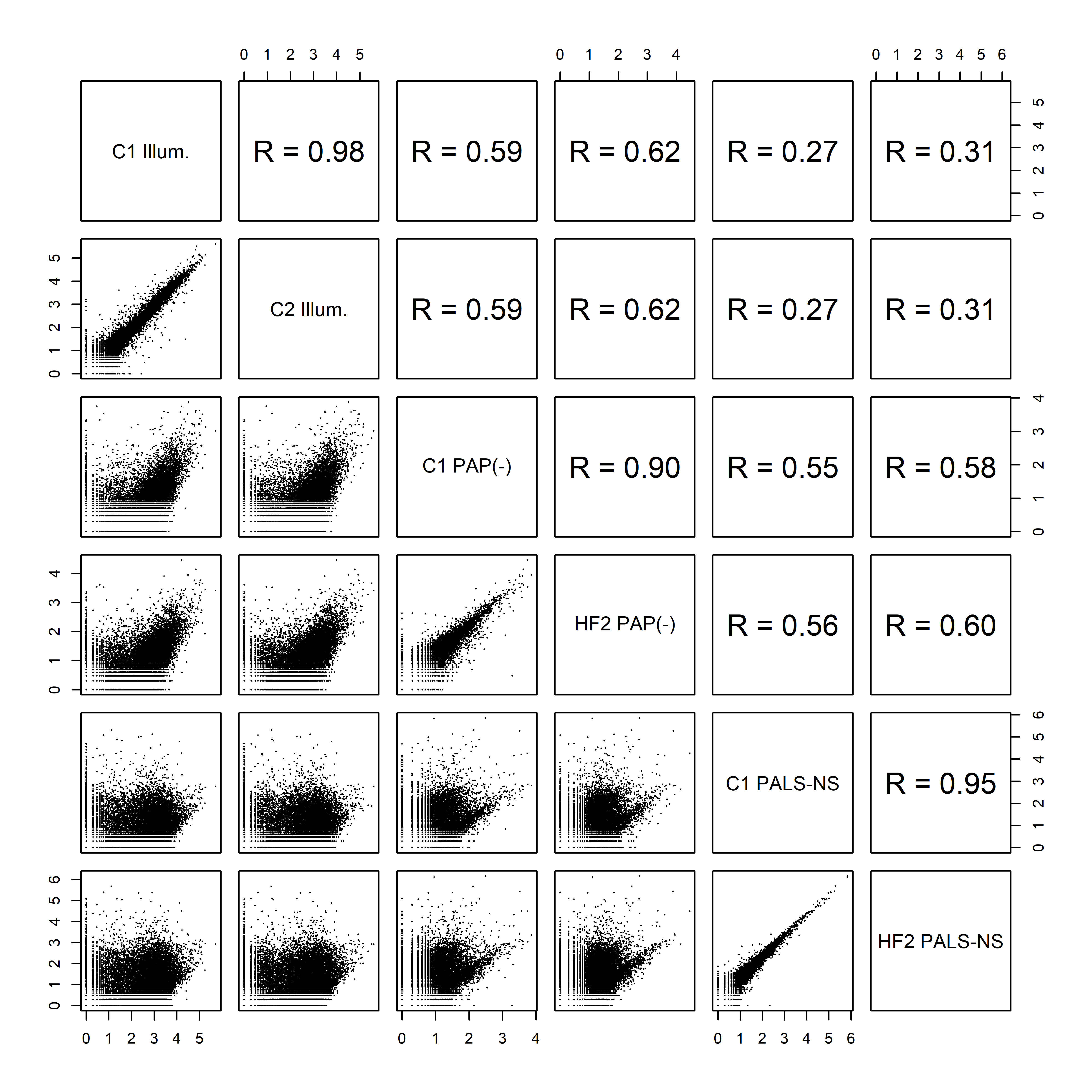

**Supplementary Figure 6** Correlation (Pearson) between raw counts of individual RNAs detected in libraries constructed from a control mouse (C1), and a mouse fed a high fructose diet (HF2) analyzed in a short RNA sequencing platform (Illumina, Illum.), the unmodified ONT long RNA protocol, PAP (-), and the PALS-NS protocol.

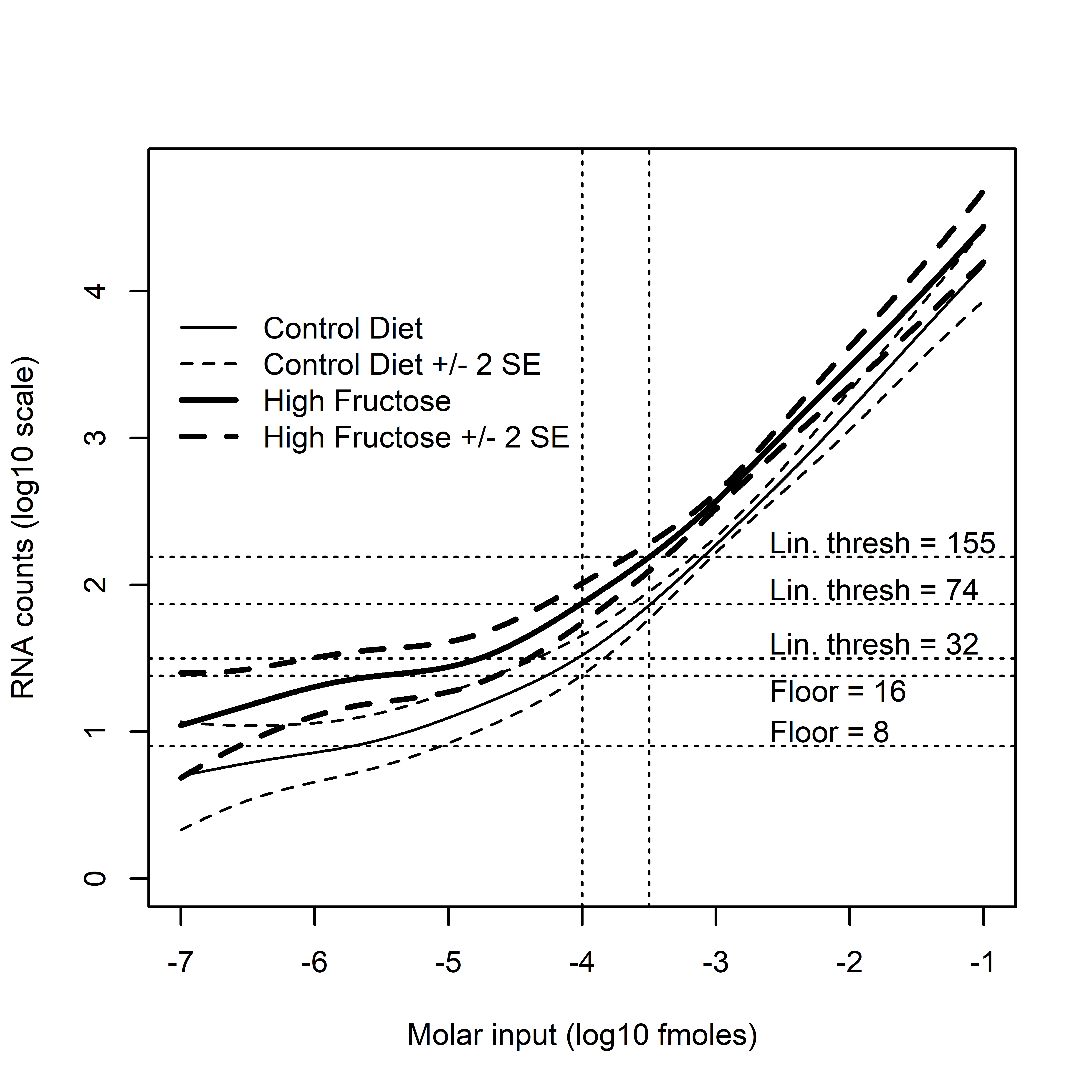

**Supplementary Figure 7** Estimated relation between RNA molar input (in log10 fmoles) in the two biological libraries, threshold of linearity and floor of detection. Two sets of thresholds were considered: a terminal set (corresponding to 32 counts in the High Fructose library and 74 in the Control Diet one) and a more proximal one (corresponding to a count of 74 in the High Fructose Library and 155 in the Control Diet one). The floor of detection, the area in which the counts don’t vary much by output was visually estimated to be at a count of 16 for the Control Diet library and 8 for the High Fructose one. Relations were estimated via Negative Binomial regression that incorporated smooth terms to adjust for molar input.
